# The proximal lipid phase of PI3K signaling is confined to the plasma membrane

**DOI:** 10.64898/2026.05.08.723799

**Authors:** Morgan M. C. Ricci, Hemani Patel, Maria J. Montoya, Kanishka L. Jayasuriya, Andrew Rectenwald, Magdalene A. K. Motter, Rachel C. Wills, Gerald R. V. Hammond

## Abstract

Class I phosphoinositide 3-kinases (PI3Ks) generate the lipid second messengers PIP_3_ and PI(3,4)P_2_ to control diverse cellular processes including growth, metabolism, and survival. Although these signals are classically thought to arise at the plasma membrane, several recent studies have proposed that PI3K signaling is propagated from intracellular membranes along the endocytic pathway. Here, we combined genomic tagging of endogenous PI3K pathway enzymes with single-molecule imaging and sensitive lipid biosensors to define the spatial organization of PI3K signaling in living cells. We find that PI3K catalytic subunits are recruited to the plasma membrane but do not undergo detectable endosomal translocation during receptor activation. Consistently, PIP_3_ and PI(3,4)P_2_ accumulation is restricted to the plasma membrane, despite enrichment of lipid phosphatases along the endocytic pathway. Functional perturbation experiments further show that degradation of PI(3,4)P_2_ occurs predominantly at the plasma membrane, indicating that both synthesis and termination of proximal lipid signals are spatially confined to this compartment. Together, these results resolve the subcellular localization of proximal PI3K signaling and support a model in which lipid second messenger production is restricted to the plasma membrane, with diversification of downstream pathway outputs occurring through redistribution of activated effector proteins rather than intracellular propagation of lipid signals.

## Introduction

Class I phosphoinositide 3-OH kinases (PI3Ks) play pivotal roles in animal physiology. They initiate a major second messenger cascade, transducing signals from receptor tyrosine kinases (RTKs) and G-protein coupled receptors. These signals activate growth, proliferation, survival, metabolism and motility in their target cells, playing major roles in organismal development and homeostasis^1^. Mechanistically, the PI3Ks phosphorylate the signature plasma membrane (PM) lipid, phosphatidylinositol 4,5-bisphosphate [PI(4,5)P_2_] at the 3-OH position, generating second messenger PIP_3_. A fraction of this PIP_3_ is then dephosphorylated to form another messenger, PI(3,4)P_2_. Both lipids then recruit and allosterically activate downstream effector proteins, such as the major serine/threonine protein kinase, AKT. Oncogenic activation of PI3K signaling occurs frequently through gain of function mutations in PI3Ks or their effectors; alternatively, the major PIP_3_ degrading enzyme, PTEN, is a major tumor suppressor. Consequently, PI3K signaling is activated in most cancers, and is currently under intense study for both pre-clinical and clinical development of small molecule inhibitors^2,3^.

Canonically, PI3K signaling occurs at the PM through recruitment of PI3Ks to activated cell surface receptors. RTK activation generates phosphorylated tyrosine residues on the receptor’s cytosolic domain, which either directly or through adaptors, engages the SH2 domains on the p85 regulatory subunit of class IA PI3K, thereby both recruiting and allosterically activating the catalytic p110 subunit^4^. However, the subcellular itinerary of PI3K signaling is potentially more complex; for example, AKT’s major downstream targets are in the cytosol, nucleus and lysosomal membranes^5^. Furthermore, the activated receptors undergo endocytosis within five minutes^6^, potentially taking PI3Ks with them^7^. Therefore, the possibility exists that PI3K signaling can occur from multiple stations of the endocytic pathway, potentially changing coupling to downstream effectors and profoundly altering signaling outcomes.

Given these varied localizations of upstream and downstream elements of the pathway, a crucial question becomes: where precisely does PI3K signaling occur in cells? An exclusive PM localization for PIP_3_ was shown by re-engineering PIP_3_-binding Pleckstrin homology (PH) domains as lipid biosensors^8–11^. However, recent evidence has shown that such sensors effectively outcompete endogenous effectors^12^, blocking signal propagation and potentially the associated traffic. Studies using organelle-localized biosensors have revealed pools of PIP_3_ and PI(3,4)P_2_ on both early endosomes and lysosomes/late endosomes (LyLE) in addition to the PM^13,14^. Moreover, PTEN has been localized to endosomal compartments^15^, and PI3Kα has even been reported to localize exclusively to microtubule-associated vesicles^16^. These studies imply activity of PI3Ks during endocytic traffic of receptors. Yet other studies reported no signaling competence in these compartments, because they lack PI(4,5)P_2_ substrate^17^. These studies relied on indirect observation of the components in fixed, processed cells or else relied on lipid-perturbing biosensors, which makes it challenging to reconcile the observations into a unified interpretation. A high resolution, real-time observation of endogenous signaling components in living cells is more likely to definitively assign localization during the dynamic evolution of signaling.

In this study, we have employed genomic tagging of the PI3K pathway’s enzymes for real-time interrogation of the pathway in living cells for the first time. Given the low copy numbers of these kinases and phosphatases, we performed sensitive, single molecule imaging to capture the spatio-temporal dynamics of the entire proximal PI3K signaling pathway. We report that PI3K activation, and resulting synthesis of both PIP_3_ and PI(3,4)P_2_ , occur exclusively at the PM. Although phosphatases degrading these lipids are indeed partially localized to endocytic compartments, all retain PM localization and loss of their activity yields enhanced lipid levels restricted to the PM. Our study thus defines the canonical PM localization as the relevant compartment for PI3K signal generation.

## Results

### PM accumulation of PI3K-generated lipids

To localize PI3K-generated lipid signals in cells, we employed genomic tagging of endogenous PI3K effector proteins AKT1 and AKT2 with mNeonGreen^12,18^, along with highly selective and sensitive genetically encoded lipid biosensors. We selected the high affinity tandem N-terminal PH domains from Myosin X (tPH) for PIP_3_^19^, and a tandem trimeric repeat of the C-terminal TAPP1 PH domain (cPHx3) for PI(3,4)P_2_^20^. Stimulation of tagged HEK293A cells with EGF revealed transient recruitment of AKT1 and AKT2 to the PM when viewed by total internal reflection fluorescence microscopy (TIRFM); peak recruitment occurred within about one minute and returned to baseline approximately five minutes later (**Fig 1A**). To detect generation of all potential cellular lipid pools, we used volumetric imaging of entire cell volumes expressing the biosensors using resonant scanning confocal microscopy.

**FIGURE 1:**
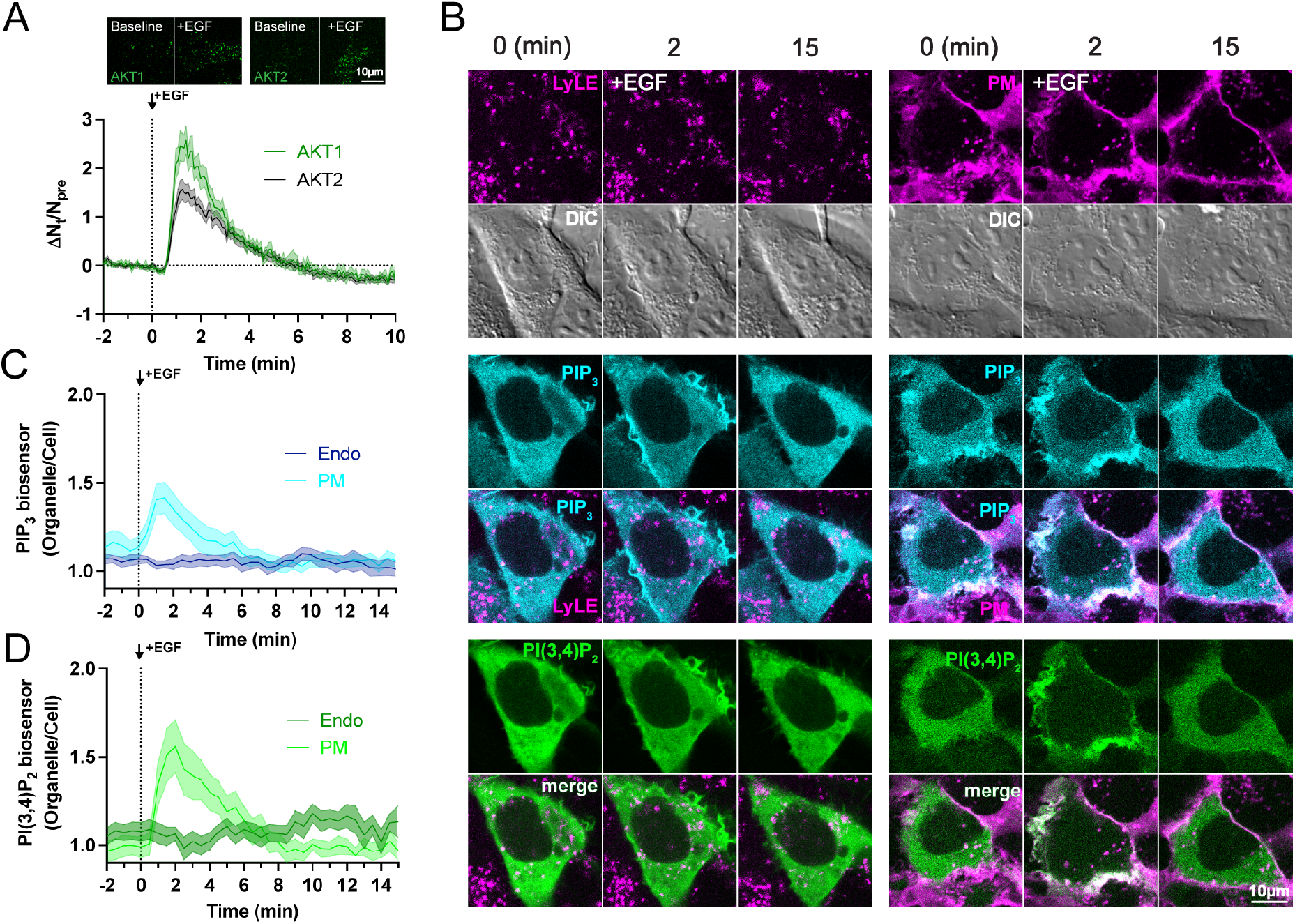
C1A-PI3K signaling initiates plasma membrane PIP_3_ and PI(3,4)P_2_ lipid synthesis. (**A**) Endogenously tagged AKT1-NG2^11^:NG^1-10^ and NG2^11^-AKT2:NG^1-10^ HEK293A cell lines treated with 10ng/mL EGF to initiate C1A-PI3K signaling after 2 min. Images show TIRF images of single tagged protein molecules recruiting to the cell surface. The graph shows the change in molecular density at the cell surface normalized to pre-stimulus baseline and are means ± s.e. of 42 AKT1-NG2^11^ and 39 NG2^11^-AKT2 cells from 16 experiments. (**B**) Representative 1.75 μm-thick confocal volumes (mean intensity projections) of cells expressing PIP_3_ biosensor tPH (cyan) and PI(3,4)P_2_ biosensor cPHx3 (green). Colocalization is shown with either PM (CellBrite; magenta) or LyLE (AF555-Dextran; magenta) at baseline (0 min), together with 2 and 15 min post-10 ng/mL EGF stimulation. (**C**) Quantification of PIP_3_ biosensor (tPH) enrichment at PM and LyLE during EGF stimulation. (**D**) Quantification of PI(3,4)P_2_ biosensor (cPHx3) enrichment at PM and LyLE. Data are means ± s.e. of 10 (PM) or 16 (LyLE) cells from 4 independent experiments.

We pre-labelled the plasma membrane with the PM-selective dye CellBrite steady 550 and labelled the endocytic pathway by a 45-minute loading with Alexa555-dextran (**Fig. 1B**). We interrogated localization of these biosensors at the markers throughout the confocal stack volume (**Figs. 1C, D**), though the images show a projection of a subset of equatorial sections for clarity (**Fig. 1B**). Stimulation with EGF revealed transient recruitment of both biosensors to the PM within one minute (**Figs. 1C, D**), with PIP_3_ returning to baseline within five minutes (**Fig. 1C**), whereas PI(3,4)P_2_ took up to seven minutes (**Fig. 1D**). However, no substantial accumulation of either was observed in endosomal membranes (**Fig. 1B-D**). Our careful volumetric analysis therefore reproduced initial observations of an exclusively PM localization of the lipids ^8–11^.

A major caveat with such genetically encoded biosensors is that they are present at concentrations matching or exceeding the PI3K-generated lipids, and in excess of the effector proteins. Therefore, they effectively out-compete these effectors and block signaling, as well as slowing the kinetics of lipid turnover^12^. This is evident from the extended accumulation of PI(3,4)P_2_ biosensor cPHx3 at the PM compared to endogenous AKT (compare **Figs 1A** and **1D**). We therefore expressed extremely low levels of biosensor to resolve single biosensor molecules in TIRFM, which accomplishes expression levels much lower than the lipids and equivalent to the effectors, preventing competition^12^. Since the effective plane of illumination in TIRFM is ∼150 nm, a region of cytoplasm above the ∼10 nm PM is captured that contains a subset of intracellular organelles, including early endosomes and LyLE (e.g.^21^ ). We expressed three biosensors; a tPH for PIP_3_, cPHx1 for PI(3,4)P_2_ and FYVE-EEA1 for PI3P as a positive control for endosomal detection^22^. The PIP_3_ and PI(3,4)P_2_ biosensors revealed diffraction limited spots with uniform intensity in TIRFM; these increased in density at the cell surface in response to EGF. On the other hand, PI3P biosensor revealed less uniform, brighter spots consistent with endosomes labelled with multiple copies of the neonGreen-FYVE-EEA1 probe (**Fig 2A**). To identify endocytic compartments, we co-transfected red fluorescent protein-conjugated marker proteins at low levels; namely Rab5a for nascent and sorting endosomes, APPL1 for nascent endosomes^23^ and clathrin light chain (CLC) for clathrin coated pits in the PM. As a control, we expressed mCherry-MAPPER, a marker of endogenous ER-PM contact sites^24^, which are distinct from clathrin coated structures, but also present as diffraction-limited spots at the PM with a similar density. Representative TIRFM images of cells expressing these probes are shown in **Fig. 2B**, with a complete set of examples for all lipid biosensors and AKT1 and AKT2 in **Fig. S1**.

**FIGURE 2:**
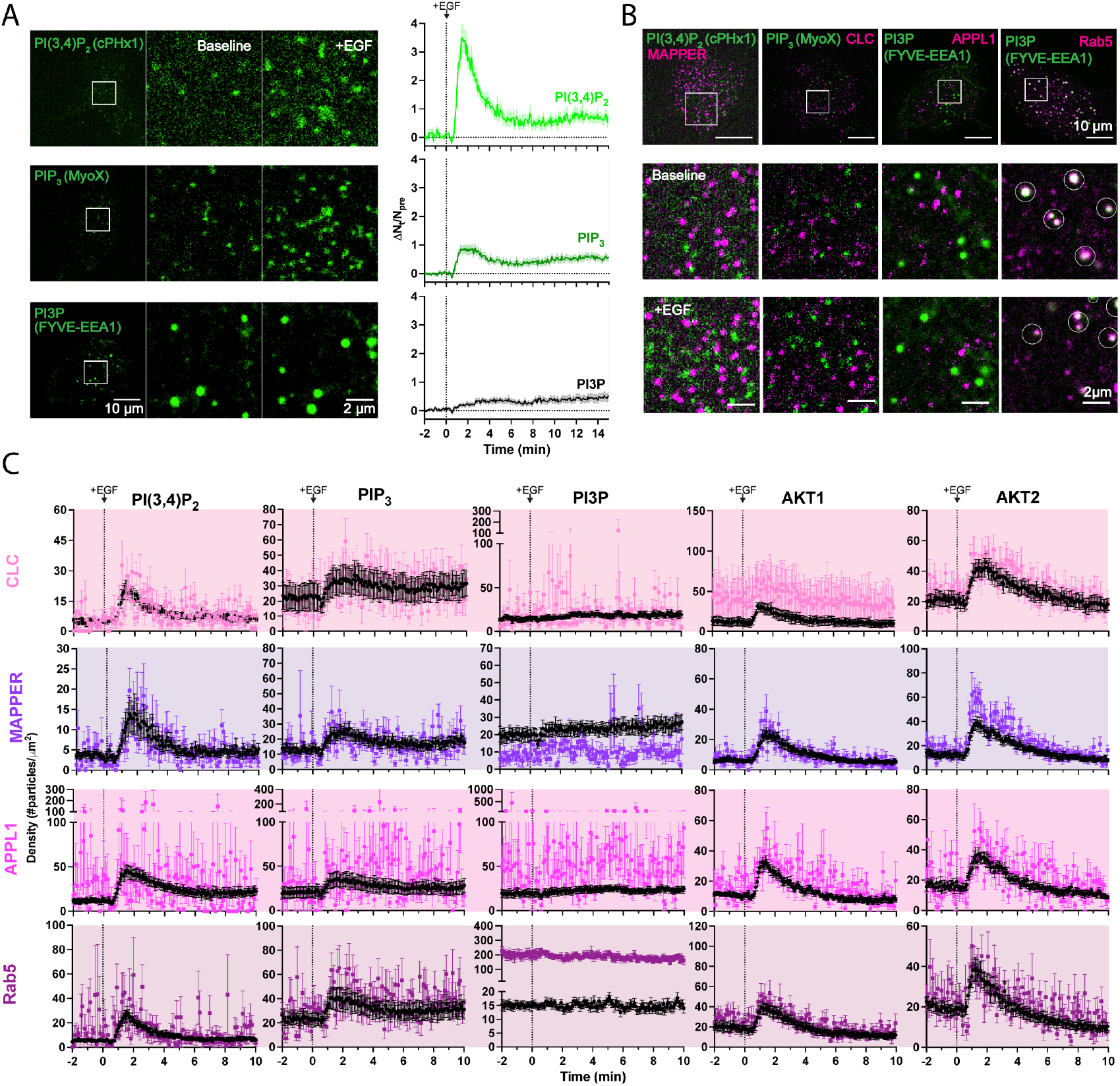
Single molecule imaging demonstrates plasma membrane restricted PIP_3_ and PI(3,4)P_2_ synthesis. (**A**) Representative TIRFM images and quantification of acutely expressed PI(3,4)P_2_, PIP_3_, and PI3P biosensors (cPHx1, tPH, and FYVE-EEA1, respectively). 10ng/mL EGF was added to cells after 2 min and kinetics of lipid synthesis quantified by the change in molecule density. (**B**) Representative images of biosensor and AKT effector colocalization with organelle markers pre- and post-EGF administration. Top row shows full cell images; middle row shows the image inset at baseline; bottom row shows the same region after EGF stimulation. PI(3,4)P_2_ and PIP_3_ biosensors do not display overlap with the marked structures, however PI3P (FYVE-EEA1) shows strong enrichment at Rab5 structures before and after stimulation (shown with white circles). (**C**) Quantification of biosensor or effector density across the whole TIRFM field or colocalized with organelle markers. Particle density was measured within a 10×10 μm region of interest. Black lines represent particle density at the cell surface; colored lines represent particle density at the organelle marker. Data are means ± s.e. of 39-42 cells from 6-8 independent experiments or 8-12 cells from 4-8 independent experiments (C).

We measured the density of biosensors (expressed in units of particles per 100 μm^2^), both across 10×10 μm TIRFM regions and co-localized with the marker proteins (**Fig. 2C**). We failed to detect enrichment of PIP_3_ or PI(3,4)P_2_ biosensor at endosomes, clathrin coated structures or ER-PM contact sites (**Fig. 2C**). Similarly, AKT generally did not show enrichment at any structure, except for a modest enrichment of AKT1 at clathrin coated structures, but no endosomes (**Fig. 2C**). Reassuringly, we spotted a robust enrichment of the PI3P biosensor at Rab5a, as expected, together with a modest enrichment at APPL1, likely reflecting nascent endosomes captured in the vicinity of larger EEA1-positive endosomes^23^. Therefore, our approach had the capability to detect endosomal PIP_3_ and PI(3,4)P_2_, yet none was evident. These lipids and their endogenous effectors AKT1 and AKT2 appeared restricted to the PM.

This approach does not suffer from the inhibitory effects seen with over-expressed biosensors, and thus confirms the majority of the PI3K-generated signals are retained in the PM. However, the necessarily low density of biosensor in this modality reduces sensitivity; in effect, a low concentration of lipid present on an endosome may be insufficient to recruit enough biosensors for detection. Therefore, to more sensitively detect the dynamics of signaling in living cells, we turned our attention to imaging the enzymes responsible for driving synthesis of these lipids: namely, the catalytic subunit of PI3K, p110.

### PI3K is recruited to and retained at the PM

Relatively few studies have directly imaged the localization of PI3Ks in living cells. Two groups have reported the insertion of a C-terminal fluorescent protein tag for either over-expression^25^ or endogenous tagging^26^, but we were discouraged from attempting this strategy, since the C-terminal “WIF” motif within p110α is essential for PM binding and efficient catalysis^27^. Therefore, addition of a ∼27 kDa fluorescent protein tag risks steric disruption of enzyme activity. Conversely, the N-terminus is in the adapter binding domain (ABD) and buried by the interaction with the p85 inter-SH2 domain^28,29^, likely causing assembly of the p85/p110 complex to be disrupted by the insertion of a globular mNeonGreen tag. That said, purification of functional, assembled p110/p85 complexes has been accomplished using N-terminal affinity tags with an 18 amino acid linker, which clears the tag from the ABD-inter SH2 interface^28,29^. We thus reasoned a similar linker could allow functional N-terminal tagging of p110 orthologues.

We designed a 16-residue flexible linker SG(GGS)_4_GG between mNeonGreen and the N-terminus of p110. AlphaFold 3 modelling^30^ of p110α and p85α reproduced the overall architecture of the complex determined by crystallography^28^, and the mNeonGreen-linker insertion was not predicted to disrupt this (**Fig. 3A**). Predicted alignment error plots showed confident prediction of the known inter-SH2 and N-SH2 domain interactions with p110α, whereas the mNeonGreen was not confidently assigned to a particular location relative to the complex, implying a desirable level of flexibility (**Fig. 3B**). We generated a plasmid encoding this p110α fusion under a truncated CMV promoter^31^ and transfected it into HEK293A cells. Low levels of p110α expression were visible as diffraction limited puncta in TIRFM, corresponding to single molecules (**Fig. 3C**). The density of these spots transiently increased after stimulating cells with EGF, indicating successful complex assembly and co-recruitment with endogenous p85. Furthermore, we could induce sustained p110α recruitment using an over-expressed FKBP-fused inter SH2 domain, driven to the PM by rapamycin-induced dimerization with PM anchored FRB^32^ (**Fig. 3C**). Collectively, these data indicated that the N-terminal mNeonGreen fusion would likely be functional in cells.

**FIGURE 3:**
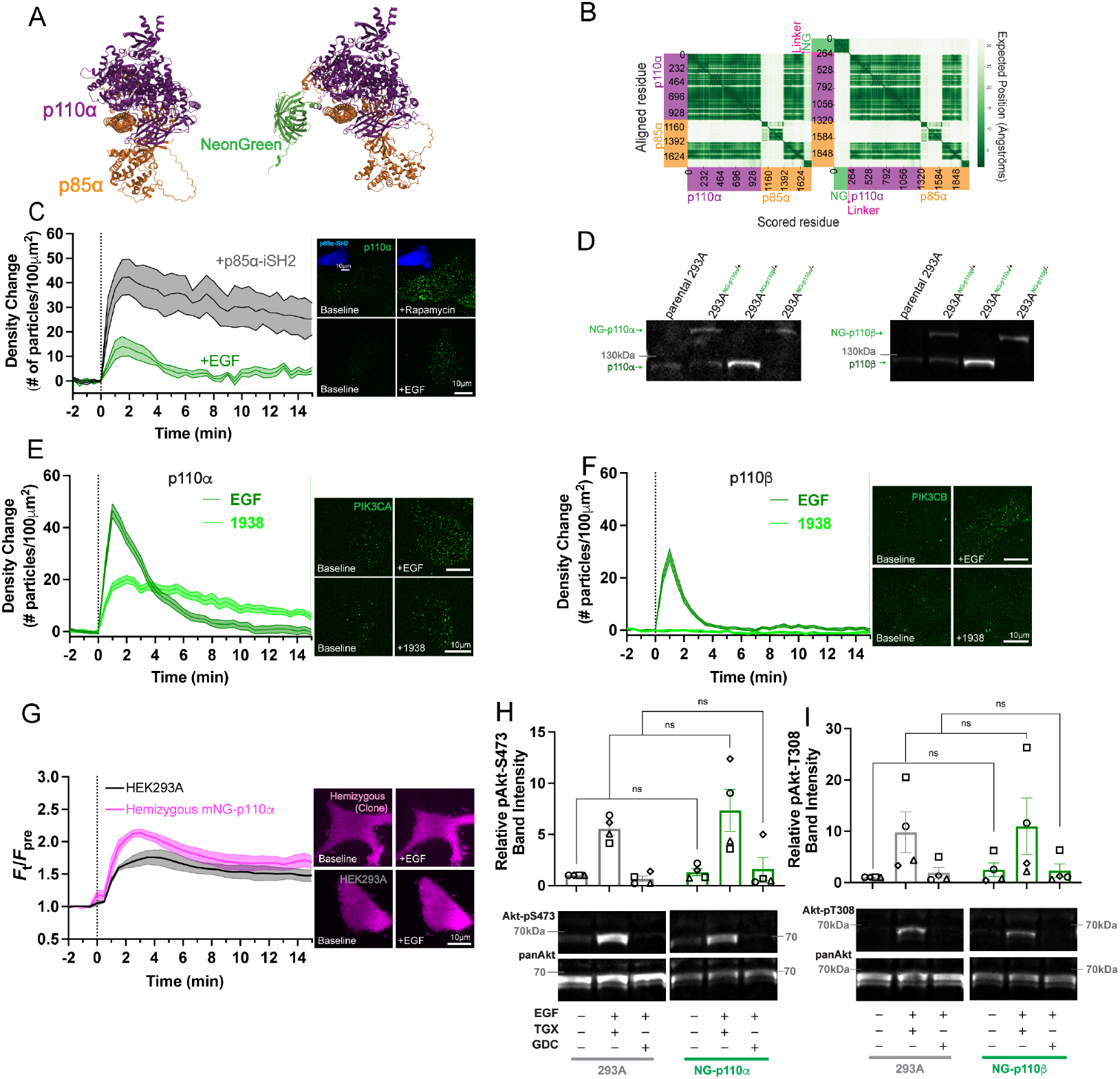
PI3K N-terminal tag with flexible linker does not disrupt enzymatic function. (**A**) Representative AlphaFold 3 structure projection of PI3Kαp110 catalytic (purple) and p85 regulatory (orange) subunit conformation, with (right) and without (left), NeonGreen - SG(GGS)_4_GG genomic insertion at the N-terminus of *PIK3CA* (p110α). (**B**) Predicted Aligned Error (PAE) heat map of PI3Kαp110-p85 prediction, with (right) and without (left)-NeonGreen-SG(GGS)_4_GG N-terminal p110αtag. (**C**) Recruitment of transiently expressed NeonGreen-SG(GGS)_4_GG-p110αin response to EGF or PM recruitment of FKBP-p85α-iSH2 with rapamycin after 2 min. Images show TIRFM images of the PM; graph shows the molecule density (grand means ± s.e. of 3 iSH2 or 4 EGF experiments with 5-10 cells each). (**D**) Western blot confirmation of NeonGreen-SG(GGS)_4_GG integration into *PIK3CA* and *PIK3CB* following CRISPR-Cas9 genomic editing. Blots of p110α and p110β are shown in un-edited parental, NG-p110α and NG-p110β heterozygous and hemizygous clones as indicated. p110α - 124,284 Da; NeonGreen-SG(GGS)_4_GG-p110α – 150,834 Da. p110β – 122,762 Da; NeonGreen-SG(GGS)_4_GG-p110β – 149,412 Da. (**E**) NG-p110α recruitment in response to 10 ng/ml or 1 μM UCL-TRO-1938. Images show TIRFM images of representative cells, and graphs are means ± s.e. of 30 (EGF) or 28 (1938) cells from three independent experiments. (**F**) NG-p110β recruitment in response to 10 ng/ml EGF or 1 μM UCL-TRO-1938. Images show TIRFM images of representative cells, and graphs are means ± s.e. of 27 cells from three independent experiments. (**G**) Quantification of PIP_3_/PI(3,4)P_2_ synthesis by Akt-PH biosensor detection in parental HEK293A and hemizygous NeonGreen-p110α cell lines upon 10ng/mL EGF + 500nM TGX-221 (TGX) p110β-specific inhibitor administration. The graph shows the grand means ± s.e. of 4 experiments (34 total cells). (**H-I**) Western blot quantification and image representation of Akt phosphorylation at the Ser473 (S473) and Thr308 (T308) activation sites in parental HEK293A and hemizygous NG-p110α cell lines. TGX and class I PI3K inhibitor GDC-0941 (GDC) were administered at 500nM and 250nM, respectively. pAkt band (S473, T308-60kDA) fluorescence was baseline corrected to the pan Akt (60kDa) loading control bands and normalized to the parental HEK293A unstimulated condition. Quantification shows no significant difference between the parental and edited lines pAkt activity. Bar graphs show means ± s.e. of 4 experiments. Results of a two-way Anova with Šídák’s multiple comparison post-hoc test (ns = P > 0.5362) are shown (six groups).

We therefore knocked this fusion into the genome of HEK293A cells using the corresponding homology directed repair template after CRISPR/Cas9-mediated targeting of the locus (see materials and methods for details). Since the copy number of p110α and p110β is estimated to be only ∼3,000^18^, HEK293A cells with integrated mNeonGreen were identified by careful imaging of post-editing cultures by fluorescence imaging in TIRF and confocal, subtracting autofluorescence to identify edited cells. The frequency of these edited cells was estimated before isolation using fluorescence activated cell sorting, yielding a polyclonal population of cells mostly heterozygous for the knock-in: these cells contained the native 110 kDa p110α band by Western blot in addition to the expected ∼110+27 kDa band (**Fig. 3D**). In parallel, we successfully generated the same fusion to a p110β allele (**Fig. 3D**). Stimulation of HEK293A cells with EGF caused transient recruitment of both mNeonGreen-p110α and mNeonGreen-p110β to the PM, again visible as an increase in the number of puncta (**Figs. 3E and F**). On the other hand, application of specific p110α activator UCL-TRO-1938^33^ caused sustained recruitment of p110α (**Fig. 3E**) but not p110β (**Fig. 3F**). Therefore, proper assembly of the tagged p110 proteins was apparent.

To verify catalytic activity, we first attempted to sub-clone the mNeonGreen-p110α cells to isolate homozygous knock-ins. However, the resulting blot showed that although the native p110 band was indeed removed, no increase in intensity of the edited mNeonGreen-p110α band was observed (**Fig. 3D**). Therefore, we believe this clone likely represents a “hemizygous” line, wherein the second allele was knocked out through non-homologous end-joining during the CRISPR/Cas9 treatment^18^. Regardless, mNeonGreen-p110α is the only copy of this protein in this cell line. Since p110α and β are the major p110 orthologues expressed in non-hematopoietic cells like HEK293A, we stimulated these cells in the presence of p110β inhibitor. Assaying PM PIP_3_/PI(3,4)P_2_ generation with the PH domain from AKT revealed no reduction in lipid synthesis in the hemizygous line relative to parental controls (**Fig. 3G**). Western blotting for AKT phosphorylation at S473 or T308 also showed no reduction in AKT activation in hemizygous cells (**Fig. 3H and I**). Therefore, our mNeonGreen fusion to p110 was functional.

Having established quantitative imaging of endogenous p110 catalytic subunits, we next sought to assign the subcellular localization of the molecules. Although TIRFM primarily illuminates the ventral PM of the cell, many intracellular organelles are also visible in the overlying layer of cytoplasm (**Fig. 2**). Given the reports of PI3K traffic through the endocytic pathway^7,13,15,16^, we tested whether the activated PI3Kα or β localized to endosomal compartments during EGF stimulation. Again, we used brief, transient transfection of endosomal markers at low levels to mark the compartments. Unlike our observations with lipid biosensors, we saw that both p110α and β showed much higher densities at clathrin coated structures after EGF stimulation, visible as the appearance of mNeonGreen puncta at mCherry-CLC spots (**Fig. 4A, B**). In contrast, no enrichment was seen coincident with the ER-PM contact site marker, MAPPER (**Fig. 4A, B** and for a full set of example images, **Fig. S2**). An enrichment with clathrin might imply internalization of the PI3Ks – yet no enrichment of either p110α or p110β was observed in complex with endosomal markers APPL1 or Rab5a (**Fig. 4A, B**). In short, we did not see evidence for endocytosis of the activated PI3Ks.

**FIGURE 4:**
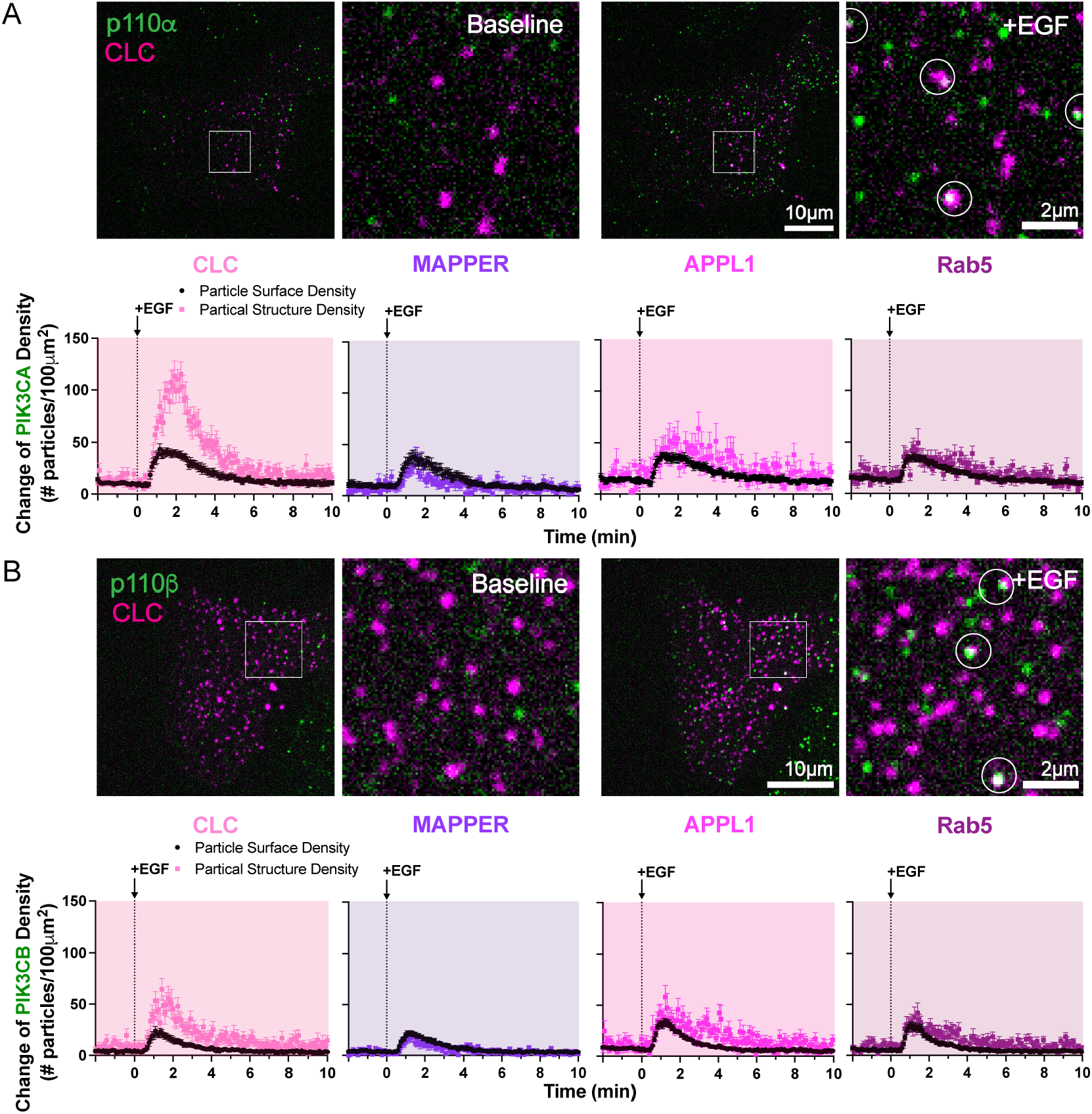
Class I PI3Ks localize to clathrin-coated structures on the PM, not to endosomes. (**A-B**) Representative TIRFM images and quantification of endogenous NeonGreen-p110α (**A**) and NG-p110β (**B**) colocalization with organelle markers. TIRFM Images of cell footprints before (baseline) and after 10ng/mL EGF administration are shown with their inset on the right. Both p110α and p110β show strong enrichment at clathrin-coated structures (CLC) upon EGF stimulation (shown in white circles). Quantification of the proteins cell surface density and colocalization with organelle markers follows the same presentation outlined in Figure 2C. Data are means ± s.e. of 8-16 cells from 6-9 independent experiments.

Why might PI3Ks be enriched at clathrin coated structures if they are not endocytosed? Previous reports have shown that clathrin coated structures in the PM can serve as signaling hubs, concentrating PI3K pathway components for efficient AKT activation without endocytosis^34,35^. We therefore reasoned that recruitment to clathrin coated structures could be required for efficient AKT activation. To test this, we knocked down clathrin heavy chain (CLTC) with small interfering RNA pools, producing robust depletion of endogenous neonGreen2-clathrin light chain (CLTA) in an edited line (**Fig. 5A**), and blocking the ability of cells to internalize Alexa647-conjugated transferrin from the cell surface (**Fig. 5B**). Despite verifying clathrin depletion in the knock-in neonGreen2-clathrin light chain cells (**Fig. 5C**), we observed no depletion of AKT phospho-S473 staining after EGF stimulation (**Fig. 5D**) by immunofluorescence. Likewise, by Western blot we observed robust depletion of clathrin heavy chain (**Fig. 5E and G**) but no depletion of AKT phospho-S473 (**Fig. 5F and G**). We also observed identical recruitment of endogenous AKT1-neonGreen2 (**Fig. 5H**) and neonGreen2-AKT1 (**Fig. 5I**), as well as undiminished synthesis of PIP_3_ and PI(3,4)P_2_ imaged with PH-AKT based lipid biosensor (**Fig. 5J**). Therefore, we conclude that clathrin coated structures are dispensable for PI3K-mediated activation of AKT. Enrichment of PI3K at clathrin coated structures is likely passive co-recruitment to these structures with the activated EGF receptors. The catalytic subunits of PI3K are not endocytosed.

**FIGURE 5:**
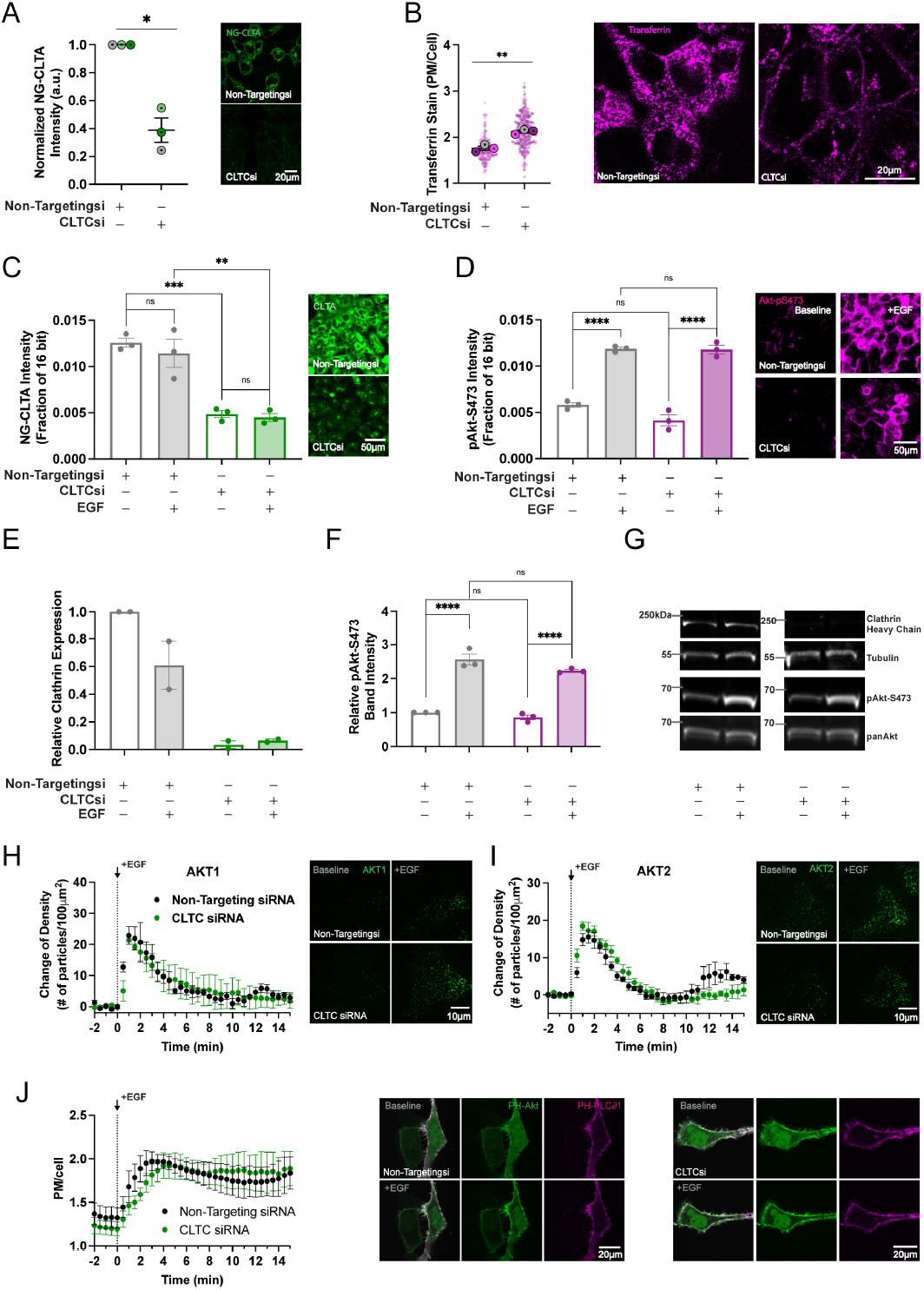
Clathrin-mediated endocytosis does not affect C1A-PI3K signaling. (**A**) Confocal micrographs of tagged NG2^1-10^:NG2^11^-CLTA HEK293A cell line treated with Non-Targeting (Neg Ctr) or CLTC-siRNA. The graph symbols show the mean fluorescence of each independent experiment (37-176 cells each); bars show grand mean ± s.e. (**B**) Uptake of Alexa647-transferrin in Non-Targeting- or CLTC – siRNA treatment. Confocal show significant Transferrin retention at the PM with CLTC -siRNA treatment. Smaller dots on the graph represent individual cell quantification and the larger symbol overlay represents the grand mean of each of the 3 experimental replicates; error bars represent the replicates mean of means ± s.e. of 163. (A-B) Results of a Welch’s t test (** = P = 0.0039, * = P = 0.02) are shown (two groups for each graph). (**C-D**) Immunofluorescence image representation and quantification of CLTA expression (C) and Akt phosphorylation at the Ser473 (pAkt-S473) activation site (D). Cells were treated with Non-Targeting- or CLTC-siRNA, then measured at baseline and 2 min post 10ng/mL EGF stimulation. Graphs show CLTA expression is reduced with CLTC-siRNA treatment, knocking down clathrin assembly, while pAkt-S473 levels reflect no difference between Non-Targeting- and CLTC-siRNA conditions. (C-D) Bar graphs data shows ± s.e. of 3 independent experiments and results of a two-way Anova with Tukey’s multiple comparison post-hoc test (C) (ns = P > 0.6698, ** = P = 0.0018, *** = P = 0.001) (D) (ns = P > 0.0644, **** = P < 0.0001). (**E-G**) Western blot quantification and representative confocal micrographs for clathrin (CLTC) expression (E) and pAkt-S473 levels (F). CLTC (180kDa) and pAkt-S473 (60kDa) bands were baseline corrected to theβ-tubulin (55kDa) and pan Akt (60kDa) loading control bands, respectively, and both were further normalized to the non-targeting-siRNA/unstimulated control condition. Data are (E) ± range of 2 experiments and (F) ± s.e. of 3 experiments and results of a two-way Anova with Tukey’s multiple comparison post-hoc test (F) (ns = P > 0.1258, **** = P < 0.0001). (**H-I**) AKT1-NG2^11^:NG^1-10^ (H) and NG2^11^-AKT2:NG^1-10^ (I) recruitment in response to 10ng/mL EGF after Non-Targeting- and CLTC-iRNA treatment. Graphs show grand means ± s.e. of 25-26 AKT1-NG2^11^ and 29-31 NG2^11^-AKT2 cells from 3 independent experiments. (**J**) Quantification and representative images of PIP_3_/PI(3,4)P_2_ synthesis by Akt-PH biosensor after Non-Targeting- and CLTC-siRNA treatments. Graphs show the grand means ± s.e. of 4 independent experiments (2-12 cells per experiment).

### PIP_3_ and PI(3,4)P_2_ are degraded in the PM

Having found clear evidence for exclusively PM-localized PIP_3_ synthesis, we next turned to lipid degradation. PI3K signal termination involves PIP_3_ breakdown via branched pathways, leading to three distinct lipid products: in the first pathway, 3-phosphorylation by PI3K is reversed by PTEN, yielding PI(4,5)P_2_^36^. In the second, inositol polyphosphate 5-OH phosphatases (INPP5) convert PIP_3_ to PI(3,4)P_2_^37,38^. This lipid is then either converted to PI4P by PTEN, or PI3P by inositol polyphosphate 4-phosphatases (INPP4)^37^. Both the major PIP_3_-selective INPP5, SHIP2, and several less selective orthologues have been localized to clathrin-coated structures in the PM^39–42^, whereas INPP4 orthologues^43–45^ and PTEN^15^ have been localized to early endosomes. It is therefore possible that PI3K lipid products may be internalized with activated receptors for endosomal signaling from endosomes, even if the PI3K does not follow. This motivated us to assess the dynamic localization of these enzymes.

We previously generated a NeonGreen2 C-terminal fusion to the major endogenous PIP_3_ 5-OH phosphatase, SHIP2^46^. For this study we also generated a C-terminal fusion of mNeonGreen to PTEN, verified by a ∼700 bp insertion with PCR (**Fig. 6A)**, and an N-terminal fusion of NeonGreen2 to INPP4A, reported by the OpenCell project^18^. Both proteins exhibited diffuse, cytosolic fluorescence when viewed in confocal which was eliminated and substantially downregulated by pooled small interfering RNA for NeonGreen2-INPP4A and PTEN-mNeonGreen, respectively (**Fig. 6B**). In TIRFM, all three cell lines yielded diffraction-limited puncta at the cell surface, albeit with differing, low density levels (≤ 12 spots per 100 μm^2^; **Fig. 6C**). Treatment of all three cell lines with EGF exhibited transient translocation of the enzymes to the cell surface, though with differing potencies: SHIP2 density almost doubled, INPP4A increased by approximately 25% and PTEN exhibited a barely 10% increase (**Fig. 6C**).

**FIGURE 6:**
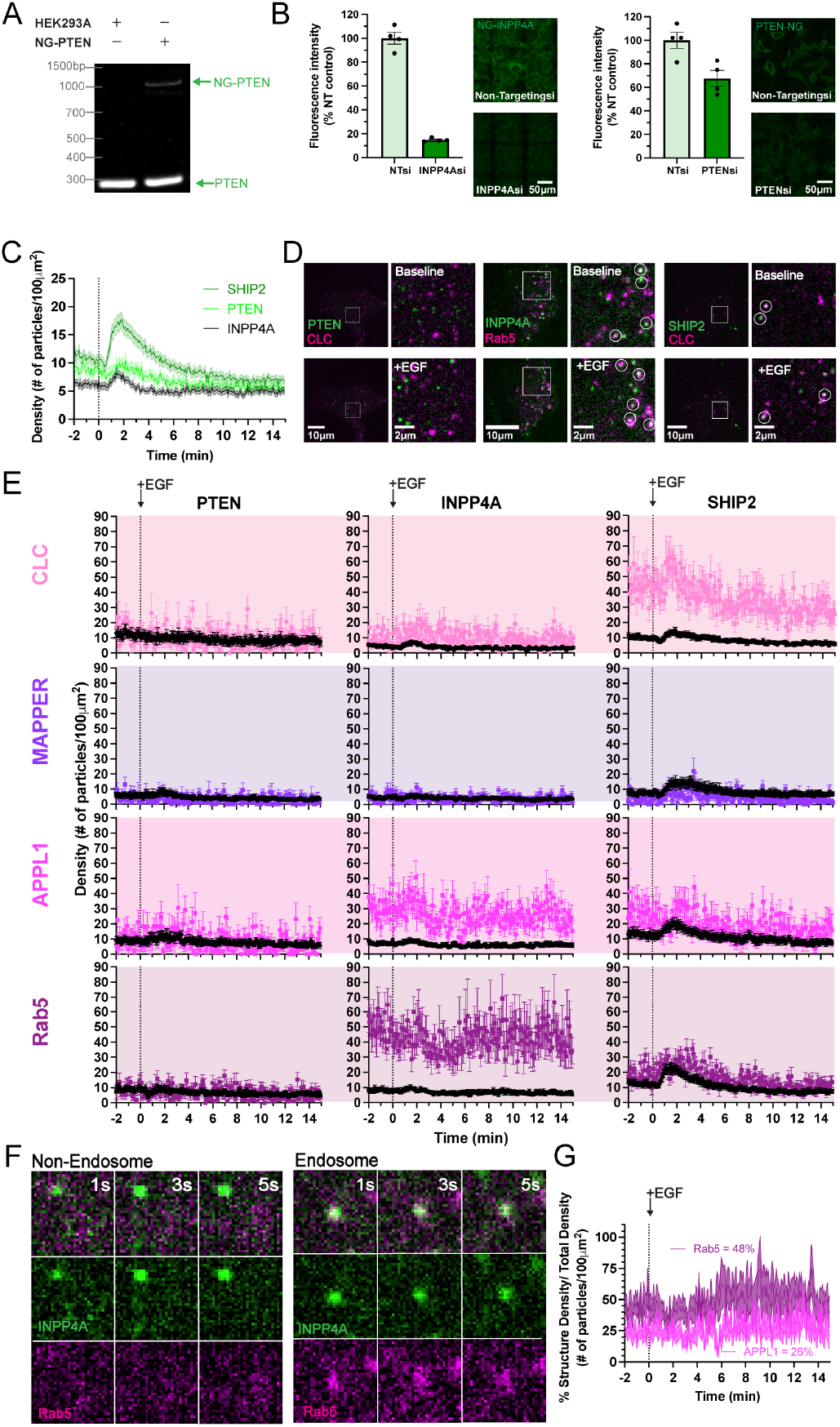
C1A-PI3K pathway endogenous phosphatases localize to cell surface. (**A**) Genotyping of PTEN-mNeonGreen (PTEN-NG) by amplification of a 997 bp fragment corresponding to the edited allele; a 280 bp fragment identified unedited PTEN expressed in both HEK293A parental and edited PTEN-NG cell lines. (**B**) Reduction of NG fluorescence by INPP4A or PTEN siRNA NG2^1-10^:NG2^11^-INPP4A (NG-INPP4A) and PTEN-mNeonGreen cell lines, respectively. Images are confocal micrographs; for quantification, autofluorescence of unedited parental HEK293A cells was set to 0 and grand means of the non-targeting control experiment was set to 100%. Data shows ± s.e. of 4 independent experiments with 57-222 cells each. (**C**) Recruitment of NG2^1-0^:NG2^11^-INPP4A, PTEN-NG, and NG2^1-10^:INPPL1-NG2^11^ in response to 10ng/mL EGF after 2 min. The graph shows the molecule density at the cell surface within a 10×10μm cell area. Data are means ± s.e. of 30-47 cells from 4-11 experiments. (**D**) Representative TIRFM images of endogenous INPP4A, PTEN and INPPL1 (SHIP2) (green) colocalization with organelle markers (magenta) pre- and post-EGF administration. Top and bottom rows show full cell images with their inset to the right; top row shows the full cell and inset at baseline; bottom row shows the same cells and insets after EGF stimulation. Co-localizing objects are shown with white circles. (**E**) Quantification of PTEN, INPP4A, and SHIP2 protein particle density at the cell surface and colocalized with organelle markers. Data are means ± s.e. of 6-16 cells from 3-5 experiments. (**F**) Image representation of single endogenous StayGold-INPP4A particle localization to non-endosomal cell surface (left) and to Rab5-labeled endosomal structure (right) at seconds 1, 3 and 5 s over a 5 second timelapse. (**G**) Quantification of percentage of INPP4A at Rab5 and APPL1 organelle markers. Graph shows means ± s.e. of 13 cells from 4 experiments.

We also tracked the co-localization of the enzymes using endocytic markers. PTEN did not show co-localization with any marker (**Figs. 6D, E**). INPP4A, on the other hand, yielded a marginal enrichment at clathrin coated structures, but a high degree of enrichment on APPL1 endosomes and markedly more so in Rab5a positive endosomes (**Figs. 6D, E**), consistent with previous reports^45^. Conversely, SHIP2 showed a dramatic enrichment at clathrin coated structures both at baseline and after EGF stimulation, but marginal enrichment on APPL1 or Rab5a positive endosomes (**Figs. 6D, E**), again consistent with previous observations^41,42^. Therefore, although we did not see evidence of intracellular PTEN stably associating with endosomes, we saw the enzymes involved in PI(3,4)P_2_ metabolism associated with the endocytic pathway. Nevertheless, careful inspection of the data revealed a more nuanced view: we frequently observed transient recruitment of StayGold-INPP4A in TIRFM that was not associated with endosomal markers, in addition to those stably associated with endosomes (**Fig. 6F**). Careful quantitative analysis of the fraction of the total TIRFM-visible neonGreen2-INPP4A revealed approximately a quarter to be in APPL1-positive and half in Rab5a-positive membranes (**Fig. 6G**). Moreover, the small but reproducible ∼25% increase in density of neonGreen2-INPP4A observed across the whole TIRFM field (**Figs. 6C, E**) was not apparent in the APPL1 and Rab5a positive membranes. It therefore appeared that in addition to the defined INPP4A endosomal pool, there is a PM-associated pool that is enriched during EGFR activation, consistent with prior reports^44^.

The PM and endocytic localization of the phosphatases means PI3K signal termination could occur at the PM, endosome or both. To delineate these possibilities, we measured accumulation of PI(3,4)P_2_ after knock-down of either PTEN, INPP4A or both. We used expression of the sensitive cPHx1 PI(3,4)P_2_ biosensor expressed at single-molecule resolvable levels for a non-invasive, sensitive measurement of lipid accumulation^12^. TIRFM imaging revealed transient recruitment of cPHx1 to the PM with EGF treatment, with no discernible effect of INPP4A siRNA (**Fig. 7A**). Conversely, the incomplete knock-down of PTEN (**Fig. 6B**) still produced a dramatic increase in the accumulated cPHx1 after EGF stimulation (**Fig. 7A**), with a subtle but detectable increase after combined knock-down of PTEN and INPP4A. In fact, the effect of PTEN siRNA was so strong that it prevented our preferred analysis method of counting the density of cPHx1 PI(3,4)P_2_ biosensor molecules^12^, since the density became too great to resolve individual molecules (**Fig. 7B**). Therefore, we had to rely on cruder measurements of background-subtracted fluorescence intensity. We therefore turned to the more conventional confocal imaging of cells expressing this cPHx1 biosensor at higher levels. EGF stimulation led to a relatively small fraction of the biosensor recruiting to the PM in control cells, producing a modest increase in the fluorescent intensity at the PM relative to the whole cell (**Fig. 7C**); this was barely perceptible in the confocal images without quantification (**Fig. 7D**). Again, knockdown of INPP4A alone had a marginal effect, whereas a substantial increase in PI(3,4)P_2_ generation at the PM was evident by PTEN knockdown (**Fig. 7C**), which was now clearly contrasted in the confocal images (**Fig. 7D**). Once again, double knockdown of both enzymes produced a subtle increase in peak PI(3,4)P_2_ generation (**Fig. 7C**). Collectively, these results show that the primary enzyme degrading PI(3,4)P_2_ under these conditions is PTEN, and that degradation of lipid by either enzyme occurs mainly in the PM. In short, the entire lipid-dependent phase of PI3K signaling appears restricted to the PM.

**FIGURE 7:**
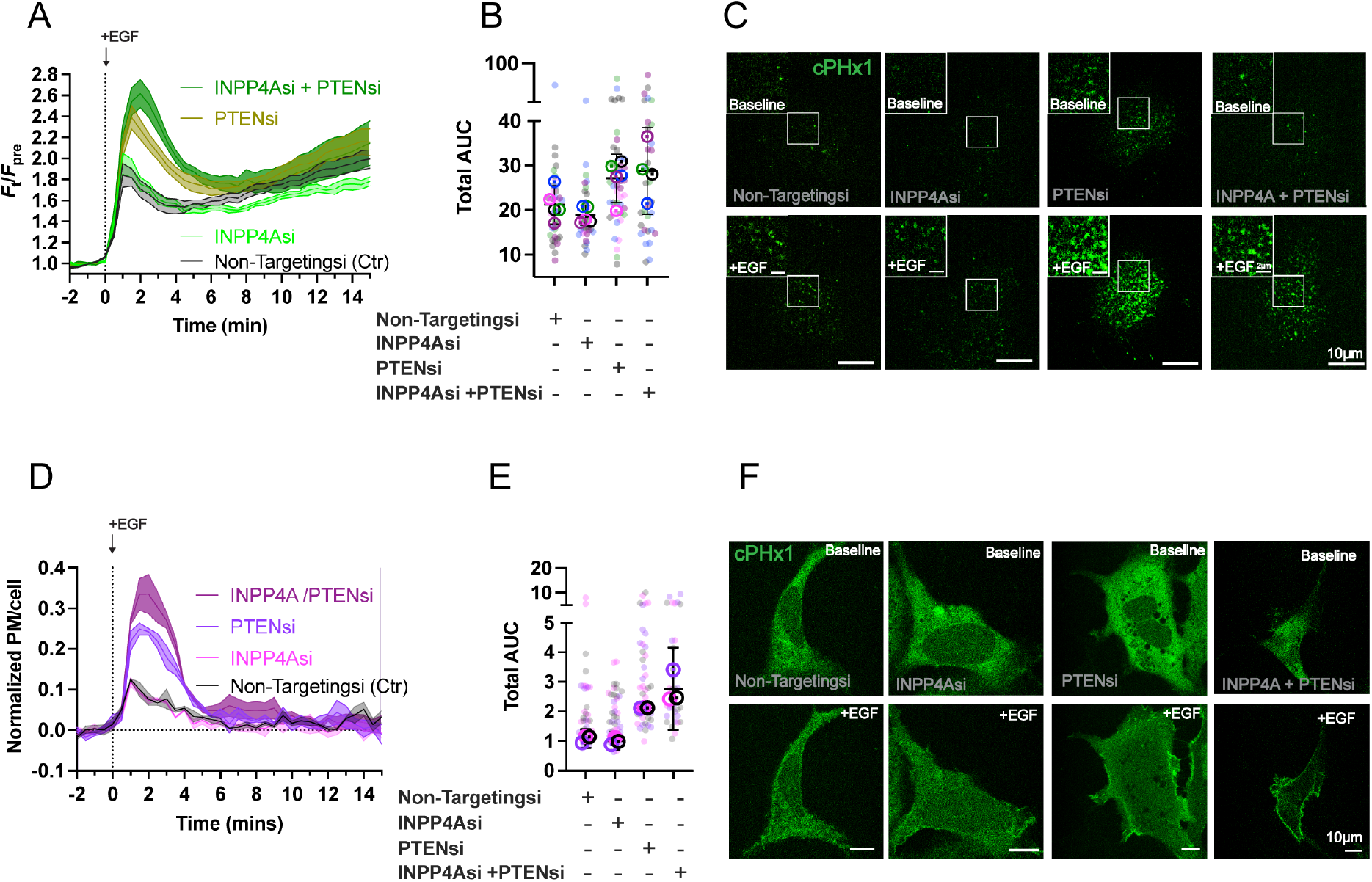
PTEN and INPP4A downregulate PI(3,4)P_2_ specifically at the PM. (**A**) Quantification of PI(3,4)P_2_ synthesis by low, single molecule resolving levels of TAPP1 cPHx1 biosensor in TIRFM during stimulation with 10 ng/ml EGF. Cells were treated with the indicated siRNA. The graph shows the grand means ± s.e. of 4-5 independent experiments (5-12 cell per experiment).**(B)** Area under the curve (AUC) quantification for each siRNA condition in (A). Individual cells are represented by small colored circles, differing in color according to specific experimental replicate. Grand means of each experimental replicate are represented with overlaying circular symbol coordinating in color to the individual cell values of the replicate. Error bars represent ± 95% confidence interval (C.I.) of the grand means ± s.e. of 3 independent experiments (10-14 cells per experiments) (**C**) Representative TIRFM images for each knockdown condition in (A) and (B). (**D**) As in (A), but quantification of PI(3,4)P_2_ synthesis was by overexpressed cPHx1 biosensor imaged in confocal. Data are grand means ± s.e. of 3 independent experiments. (E) AUC quantification of each siRNA condition in (D) as described for (B). (**F**) Representative confocal images of cPHx1 in each knockdown condition in (D) and (E). Data are means ± s.e. of 33-45 cells from 5 independent experiments (A-B) or 33-42 cells from 3 independent experiments (D-E).

## Discussion

PI3K’s central role in physiology has been elucidated by a combination of highly innovative genetic and physiological experiments that defined biological function, together with elegant and meticulous biochemical and structural characterization of the enzymes’ regulation^2,4^. However, the cell biological basis of this regulation – when and where the enzymes and ipids localize during signal propagation – has been much less clearly defined. Here, we used a combination of exquisitely sensitive lipid biosensors and high-resolution imaging of endogenous enzymes to define the spatial regulation of PI3K signaling in living cells with single-molecule resolution. Our data are consistent with the canonical model that PI3K activation and lipid signaling occurs exclusively at the plasma membrane. The lipid second messengers do not appear to be synthesized nor signal from intracellular compartments.

A PM-exclusive localization for PI3K-derived second messengers likely facilitates the ability to precisely modulate the signal. The precise dynamics of PI3K signals shape the pathway’s ultimate output; variations in amplitude, duration and frequency of PIP_3_ generation can all be decoded by downstream effectors to transduce different physiological outcomes^47,48^. By restricting the lipid signals to the PM, the pathway is primed at the ultimate source of its substrate, PI(4,5)P_2_, and the receptors that acutely respond to the extracellular cues that trigger them. Such proximity is likely crucial for precise modulation of signal dynamics. Alternatively, PI3K signals propagating through the endocytic pathway would accrue significant kinetic delays, limiting modulation to the tens of minutes associated with vesicular traffic.

Curiously, endocytic traffic of PI3K-generated lipids has been implicated after their detection in the early and late endocytic pathway by organelle-specific biosensors^13^. In this case, the kinetics of lipid generation only lagged slightly behind the plasma membrane, and occurred much faster than the traffic of the activating receptors. Endocytic traffic of PIP_3_ and PI(3,4)P_2_ would be especially surprising, given the gauntlet of clathrin-coated structure and endosome-associated phosphatases that dephosphorylate these lipids, as shown here (**Fig. 6**) and in prior studies, e.g.^41,42,44,49^. In this case, we suspect that rapid non-vesicular lipid transport, such as via the PIP_3_ binding TIPE3 protein^50^ , could play a role. In that case kinetic control of the signal dynamics would still be under ultimate control of the plasma membrane-localized enzymes, with only a minor lag. This model would therefore be largely compatible with our observations here.

Another consequence of restricted signal generation in the plasma membrane would be to set up a powerful coincidence detection mechanism. This could disambiguate receptor-derived second messengers at the PM from the effects of the same lipid molecule elsewhere in the cell. A notable example is the generation of PI(3,4)P_2_ from PI4P at LyLE by the β orthologue of class 2 PI3K. In this context, the lipid represses mTORC1 activation at the LyLE^51^ , as opposed to plasma membrane-derived class IA PI3K signals, which activate it.

Whilst our studies were underway, there was a similar report of endogenous tagging of p110 subunits^26^. Again, a similarly long nine or fifteen amino acid linker was used, in this case between the C-terminus and the fluorescent protein tag. As with our N-terminal tag, this likely allows the tag to clear the C-terminus and prevent steric hinderance of membrane interaction. Clear EGF-induced PM translocation of p110α and β was also observed, though the localization to stations of the endocytic pathway were not interrogated. Instead, clear recruitment to focal adhesions was reported in otherwise unstimulated cells. This may well derive from focal adhesion kinase mediated Src activation^52^ , which can in turn activate PI3K via adapter proteins such as PLEKHS1^53^. Nonetheless, the data reported in that study are entirely consistent with the plasma membrane restricted signaling that we report here.

Our data implicated PTEN as by far the most important enzyme for degrading PI(3,4)P_2_ in HEK293A cells (**Fig. 7**). This contrasts with a previous report, where loss of both PTEN and INPP4B were required to obtain substantial PI(3,4)P_2_ accumulation in MCF10a breast epithelial cells^37^. The difference likely lies in the enzyme complement of these cells. INPP4B is not detectable in HEK293A cells^18^, and PTEN is present in the plasma membrane of these cells at substantially higher concentrations than INPP4A. It’s therefore likely that MCF10a have a higher concentration of constitutively plasma membrane localized INPP4 activity in the form of INPP4B. These differences aside, MCF10a cells were also observed to have PI(3,4)P_2_ accumulated in the plasma membrane^37^.

If the majority of PI3K-derived lipid second messengers are dephosphorylated in the plasma membrane, why are the phosphatases also localized along the endocytic pathway? An obvious reason is to prevent “leak” of the lipids into the endocytic pathway. Indeed, clathrin mediated endocytosis has been found to have a very limited ability to sort lipids^54^, which are likely to passively enter by diffusion. In addition, clathrin mediated endocytosis requires depletion of PI(4,5)P_2_ for uncoating, and the INPP5 enzymes utilized for this task all exhibit potent activity against PIP_3_^38^. It follows that any PIP_3_ diffusing into the nascent vesicle is doomed for conversion into PI(3,4)P_2_. This would join the small pool of PI(3,4)P_2_ generated during vesicle scission by class 2α PI3K during constitutive clathrin mediated endocytosis^55^, which explains the requirement for INPP4A in endosomes^45^. However, the constitutive pool of PI(3,4)P_2_ associated with endocytosis is vanishingly small compared to that generated by the class IA PI3K pathway in the plasma membrane^37^, and not detected with even the most sensitive biosensors^20,55^.

The definitive localization of proximal PI3K signaling to the plasma membrane poses important questions going forward: PIP_3_ and PI(3,4)P_2_ may be largely restricted to the plasma membrane, but the targets of their effector proteins are certainly not. For example, the major PIP_3_/PI(3,4)P_2_ effector AKT relies on the lipids both for membrane recruitment and allosteric activation^56,57^. However, it remains unclear whether lipid engagement must be continuous after the proteins become phosphorylated^58^. This is a central question, since AKT substrates are in the cytosol, LyLE and nucleus^5^ , yet we observe exclusive AKT recruitment at the plasma membrane. We can envisage two possibilities: AKT, once phosphorylated, can disengage from the membrane yet retain sufficient activity to phosphorylate its targets after diffusing through the cytosol. Alternatively, if sustained lipid binding is required for activity, downstream signaling may instead depend on transient encounters between substrates and activated AKT at the plasma membrane. In either case, our findings place the plasma membrane as the obligate site of signal initiation and early propagation in the PI3K pathway, implying that spatial regulation downstream of lipid generation must occur through redistribution of activated effectors rather than redistribution of the lipid signals themselves. Resolving how this transition from membrane-confined lipid signaling to distributed substrate phosphorylation occurs will be central to understanding how PI3K signaling achieves both specificity and versatility in physiological contexts. This will be especially important in the continued pre-clinical efforts to block specific pathological impacts of PI3K activation, while sparing physiologically crucial processes.

## Materials and Methods

### Cell culture, Transfection, Gene Editing

HEK293A cells (ThermoFisher, R70507) were kept in humidified incubators at 37°C, 10% atmospheric CO_2_, and cultured in low glucose Dulbecco’s Modified Eagle Medium (DMEM), GlutaMAX™ supplement, pyruvate (ThermoFisher Scientific,10567022) growth medium supplemented with ThermoFisher Scientific 100 μg/mL streptomycin + 100 U/mL penicillin (15140122), 10% heat-inactivated fetal bovine serum (10438-034), and 0.1% (vol/vol) chemically defined lipid supplement (11905031). The cells were routinely screened for mycoplasma and passaged 1:5 by rinsing in Ca/Mg-free phosphate buffered saline and dissociation in Trypsin-like enzyme, before diluting in medium and splitting into fresh vessels.

Cells were seeded in 6-well plates for both western blot and immunofluorescence protocols. Though the cells adhere to the plastic well bottoms sufficiently for western blot experiments, the 40 mm #1.5 glass coverslips (ThermoFisher Scientific, 50-153-2238) placed in the wells for immunofluorescence and the 35-mm circumferential, 20mm #1.5 optical glass bottom dishes (CellVis, D35-20-1.5-N) used in microscopy, were pre-coated for 30 min at 37°C with 10-20μg ECL cell attachment matrix (Millipore Sigma, 08-110).

Transfection of the 6-(35mm) well plates and 35mm glass bottom dishes were performed with ∼1μg DNA precomplexed to 3μg Lipofectamine™ 2000 in 0.2mL Opti-MEM™ | Reduced Serum Medium, GlutaMAX™ Supplement (ThermoFisher Scientific, 11668019). For overexpression of constructs, the pre-complexed DNA solution was added to cells and incubated for 4 hours. Following this incubation time the solution was removed and the cells serum-starved 24-hr prior to stimulation in serum-free Gibco™ FluoroBrite™ (ThermoFisher Scientific, A1896702) with 0.1% chemically defined lipid concentrate (ThermoFisher Scientific, 11905031), 0.1% Gibco™ Bovine Albumin Fraction V (7.5% Solution) (ThermoFisher Scientific, 50-121-5315) and 25 mM NaHEPES, pH 7.4. For single molecule-level expression, the pre-complexing steps are unchanged, however the DNA complex solution is added to the cells ∼2hr prior to stimulation and subsequent measurement. The cells are still serum-starved 24 hr prior to stimulation, however transfection occurs same day.

The employed CRISPR/Cas9-mediated gene editing approach follows previously illustrated protocols^59,60^. The major PI3K pathway signaling components were selectively targeted with CRISPR/Cas9 machinery and homology-directed repair to integrate the mNeonGreen tag into specific loci. Two main tagging approaches were utilized: in the first, full mNeonGreen coding sequence is inserted into the gene of interest; in the second, a split mNeonGreen variant (NG2^11^) is incorporated in HEK cells expressing the rest of the neonGreen2 protein (referred to as HEK293^NG2-1-10^ cells)^60,61^. A third tagging strategy was used to incorporate fluorescent StayGold into INPP4A, employing the same genomic editing approach utilized for tagging with mNeonGreen.

The guide RNA protospacer sequences and homology-directed repair templates were introduced to the cells by a Neon electroporation system. The homology-directed repair templates were synthesized by IDT as a single-stranded and double-stranded “Alt-R®” sequences for NG2^11^ and full mNeonGreen genomic integration, respectively. Each template followed a general structure containing homology arms for gene alignment, a flexible linker between the gene’s ORF and that of mNeonGreen, and the NG2 or NG2^11^ insertion sequence. As previously described^12^ , 10 pmol of gRNA and 10pmol of TruCUT™ Cas9 protein V2 (ThermoFisher Scientific, cat#A36498) were preincubated together for 20 min in 4.2-5μL buffer R, followed by the addition of 100 pmol of homology-directed repair template. The solution was mixed and subsequently combined with 200,000 suspended cells in 5μL buffer R. Electroporation used a single 20-ms, 1,500V electroporation pulse. Following electroporation, cells were placed in antibiotic(Pen/Strep)-free DMEM and incubated for 12-16 hours at 37°C, 10% atmospheric CO_2_. For some of the experiments, 1.7μL of Alt-R™ HDR Enhancer V2 (IDT, 10007921) was added to the media during incubation. The antibiotic-free media was then subsequently replaced with complete DMEM and the cells cultured and screened for mNeonGreen fluorescence by confocal microscopy 48-hours post-seeding. With ≥1% edited cell population, fluorescence-activated cell sorting (FACS-University of Pittsburgh Flow Cytometry Core) was employed to isolate NG2 or NG2^11^ positive polyclonal populations. Further details on the templates used, genomic location of fluorescent tag and the specific proteins tagged are provided in Key Resources Table.

Full length mNeonGreen was inserted into the PTEN gene at the carboxy terminus. For confirmation of this successful integration, genomic DNA was extracted by the Purelink™ Genomic DNA Mini Kit. This purified DNA was diluted to 10ng/μl and the insertion region amplified with polymerase chain reaction. The resulting sample was run on a 2% agarose gel; a 997bp product confirmed the ∼700bp mNeonGreen integration while a 280bp product reflected the native un-tagged protein.

### Chemicals

Corning™ Epidermal Growth Factor (ThermoFisher Scientific, CB-40052) was resuspended in Mili-Q water to 0.1mg/mL and diluted to 10ng/mL in imaging medium for cell stimulation. Stocks were stored at -20°C for extended time and 4°C when actively used post-thaw (EGF was not subject to repeated freeze-thaw cycles). Rapamycin (Millipore Sigma, 553210-100UG) was dissolved to 1mM with DMSO and diluted to 1μM for cell stimulation. Stocks were stored at -20°C. PI3Kα activator (UCL-TRO-1938TM; CancerTools.org) was resuspended to 10mM with DMSO and diluted to 1μM for cell stimulation. Stocks were stored long term at -80°C and short term (∼3 months) at -20°C. PI3Kβ inhibitor TGX-221 (MedChemExpress, HY-10114) was dissolved in DMSO to 10mM and diluted to 500nM for cell exposure; stocks were stored at -20°C. Class I PI3K inhibitor GDC-0941 (Millipore Sigma, 5.09226) was resuspended in DMSO to 2mM and diluted to 250 nM for cell exposure; stocks were stored at -20°C.

To label the plasma membrane and endosome structures in Figure 1-z-stack A1R imaging experiments, CellBrite^®^ Steady 550 (Biotium, 30107) and Dextran Alexa Fluor™ 555 (ThermoFisher Scientific, D34679) were used, respectively. CellBrite^®^ Steady 550 was incubated overnight with enhancer, both concentrated 1:1×10^5^ in SF-CHIM+0.1%BSA imaging solution. To label endosomes, cells were loaded with Dextran Alexa Fluor™ 555 on the same-day as imaging, diluted 1:1000 in SF-CHIM+0.1%BSA. The cells were incubated with the Dextran stain for 45 min at 37°C. Both the CellBrite^®^ and Dextran stains were performed with cells transfected with a HaloTag-fused lipid biosensor, therefore Janelia Fluor 646 HaloTag Ligand mix (Promega, GA1120) was added to the dishes at 1nM concentration and incubated for 15min at 37°C). Each dish was then washed three times with phosphate-buffered saline (PBS) and incubated for 30min in SF-CHIM+0.1%BSA and subsequently imaged.

Transferrin Alexa Fluor™ 647 Conjugate (ThermoFisher Scientific, T23366) employed in Figure 5 was stored at 5mg/mL and incubated on cells at 25μg/mL made up in SF-CHIM+0.1%BSA for 15min at 37°C. To label the plasma membrane, the solution was replaced with CellMask™ Orange stain (ThermoFisher Scientific, C10045) at 2.5ng/μL also made up in the SF-CHIM+0.1%BSA imaging media and directly applied to the glass well in the center of the dish. The dish was then incubated at 37°C for an additional 5min. Following stimulation with Transferrin and staining with CellMask™ Orange, the cells were washed three times with 1mL pre-warmed SF-CHIM+0.1%BSA and finally imaged in 2mL SF-CHIM+0.1%BSA media.

### Western Blotting

To measure phosphorylated Akt levels, cells were cultured in 6-well plates at 25% confluency. 10μL of 5μM siRNA working stock (1-20μM stock: 4-RNase-Free water) was diluted in 190uL SF-CHIM and 6.6uL DharmaFECT (horizon, T-2001-01) was diluted in 193.4μL, both mixtures sat for 5min then were combined and incubated at room temperature for 20min. Following incubation, experiments testing specific knockdown effect on Akt phosphorylation levels would then have the 6-well plates media replaced with siRNA solution added to 1600uL antibiotic(Pen/Strep)-free DMEM. Cells were incubated for 24 hr at 37°C, then replaced with fresh siRNA solution and incubated for an additional 4 hr. The second siRNA solution was then removed and the cells incubated for 24 hr with regular Complete DMEM. After 24hr, the media was removed and the cells incubated overnight with SF-CHIM +0.1%BSA. Cells were stimulated with EGF 3 days following initial siRNA exposure for 2min then lysed in RIPA buffer (ThermoFisher Scientific, 89900) supplemented with protease (Millipore Sigma, 539131-1VL) and phosphatase (Millipore Sigma, 524627) inhibitor cocktails. Following 15 min incubation on ice, the lysed cells were scraped and centrifuged at 13,000rpm for 10min. 40μL of this sample was mixed with 15μL of 4x LDS (Bolt™ LDS Sample Buffer; ThermoFisher Scientific, B0008) and 6μL reducing agent (10x Bolt™ Sample Reducing Agent; ThermoFisher Scientific, B0009) then boiled at 100°C for 5min and cooled >2min on ice. Samples and ladder marker (PageRuler Prestained NIR Protein Ladder; ThermoFisher Scientific, 26635) were then loaded in precast polyacrylamide gels (Bolt™ Bis-Tris Plus; ThermoFisher Scientific, NW04122BOX, NW00082BOX, NW00102BOX) set in 1xBolt MOPS Running Buffer (ThermoFisher Scientific, B0001) and run at 165V for 45min. The gel was then isolated and transferred onto a Nitrocellulose membrane (ThermoFisher Scientific, LC2000) at 10V for 1hr. The resulting membrane was blocked for 1hr at room temperature with 1XTBST (Tris Buffered Saline; ThermoFisher Scientific, BP24711 and 0.05% Tween; Millipore Sigma, P1379) containing 10% Milk (Cell Signaling Technology; 9999S) or 1XTBST containing 5% BSA (Gibco™ Bovine Albumin Fraction V; ThermoFisher Scientific, 50-121-5315)) when staining for Phospho-Akt (Ser473) (D9E) rabbit monoclonal antibody (Cell Signaling Technology, 4060) or Phospho-Akt (Thr308) rabbit monoclonal antibody (Cell Signaling Technology, 13038), respectively. Blots measuring phospho-Akt levels were stained with primary antibodies, Phospho-Akt (Ser473), Phospho-Akt (Thr308), Akt (pan) mouse monoclonal (Cell Signaling Technology, 2920), and secondary antibodies Goat anti-Rabbit IgG (H+L) Highly Cross-Adsorbed Secondary Antibody, Alexa Fluor™ 647 polyclonal (ThermoFisher Scientific, A-21245) and Goat anti-Mouse IgG (H+L) Highly Cross-Adsorbed Secondary Antibody, Alexa Fluor™ Plus 800 (ThermoFisher Scientific, A32730). In parallel to Akt-pS473, clathrin heavy chain (CLTC) was also stained for in CLTC knock-down conditions with primary Anti-Clathrin heavy chain mouse monoclonal [X22] (Abcam, 2731) and Beta-Tubulin (9F3) rabbit monoclonal (Cell Signaling Technology, 2128) antibodies.

Blot images were captured in the Alexa647 and DyLight800 channels. Membranes were stained so that one channel detected the bands of the loading control and the other of the target measurement. The representative TIF images generated were imported into FIJI and background corrected with a rolling ball radius of 30.0 pixels to remove uneven backgrounds (note: 30 pixels is larger than the thickest bands). Regions of interest (ROIs) were then drawn with the Rectangle tool around each band and a portion of blank gel space in both channels to set as background. The ROIs intensities were then measured and copied into Graphpad PRISM.

With Graphpad PRISM, each channels band intensities were baseline-corrected to their specific background ROI. The baseline-corrected intensities for the loading control bands were then subtracted from those of the blots target measurement. The resulting subtraction values were normalized to the standard of the target measurement; standards included the parental HEK293A cells in the unstimulated condition for pAkt-S473/T308 blotting and the non-targeting-siRNA, unstimulated control setting for Clathrin-knockdown blotting.

### Immunofluorescence

Cells were seeded onto glass coverslips and cultured in 6-well plates. Following siRNA treatment and stimulation, the cells were fixed using 4% formaldehyde in 10xPBS for 15 min at room temperature. After 15 min the coverslip was washed with 50mM ammonium chloride in 1xPBS solution three times, then incubated for 30 min in 1XPBS blocking solution supplemented with 0.2% TX-100 (Triton™ X-100; Millipore Sigma, TX1568-1) and 5% NGS (normal goat serum; ThermoFisher Scientific, PCN5000). The coverslip was then stained for 1hr with 1:400 rabbit monoclonal Phospho-Akt (Ser473) (Cell Signaling Technology, 4060) and 1:1000 mouse monoclonal Anti-Clathrin heavy chain (Abcam, 2731) primary antibodies in block solution. Following a wash with 1xPBS, the coverslip was then stained for 30 min with block solution supplemented with 1:400 Goat anti-Rabbit IgG (H+L) Highly Cross-Adsorbed Secondary Antibody, Alexa Fluor™ 647 polyclonal (ThermoFisher Scientific, A-21245) and polyclonal Goat anti-mouse IgG1 Cross-Adsorbed Secondary Antibody, Alexa Fluor™ 488 (ThermoFisher Scientific, A-21121) secondary antibodies, 1:40,000 CellMask™ Orange stain (ThermoFisher Scientific, C10045), and 1:500 DAPI (4’-6-Diamidino-2-Phenylindole, Dihydrochloride; ThermoFisher Scientific; D1306). Following fixation and staining, the coverslip was washed with 1xPBS and Millipore water, then left to dry. Using ProLong™ Diamond Antifade Mountant (ThermoFisher Scientific, P36961), the coverslip was mounted to a larger rectangular coverslip and imaged on a Nikon A1R confocal microscope. The cells were captured as a large field image using the 20X plan apochromatic 0.75 NA air immersion objective with an open pinhole at 255.43 μm. Excitation was directed by a LU-NV laser combiner using the 405.0 nm line to collect DAPI emission using a 425-475-nm filter, 488.0 nm to collect Alexa488 emission at 500-550-nm, 561.0 nm to collect CellMask™ Orange at 570-620 nm, and 647.0 nm to collect Alexa647 at 650-850 nm. Identification and analysis of individual cells followed the same procedure outlined previously in Holmes et al 2025^12^, where the “Huang” method, Watershed function, and Voronoi function allowed for appropriate cell segmentation and defined ROIs^12^. ImageJ was further utilized to measure the raw 12-bit intensity of the AKT-pS473 and clathrin heavy chain channels within the ROIs^12^.

### Microscopy

Confocal imaging was conducted by a Nikon A1R microscope composed of a Nikon TiE inverted stand attached to an A1R resonant scan head and a heated stage incubator set to 37°C (Tokai-Hit). LU-NV laser combiner with four-line excitation of 405-, 488-, 561-, and 640-nm allowed collection of blue, green, yellow/orange, and far/infrared channels along with transmitted light for DIC. 100X 1.45 NA plan-apochromatic oil-immersion objective and 8 to 16 frame averages were used.

TIRFM imaging was conducted by another Nikon microscope composed of a Nikon TiE inverted stand fitted with a TIRF illuminator arm fiber linked to a four-line Oxxius L4C wavelength combiner and a heated stage incubator set to 37°C (Tokai-Hit). Images were acquired through a 100X 1.45 NA plan-achromatic oil-immersion objective by a Hamamatsu Fusion-BT sCMOS camera. Imaging of excitation wavelength 405-nm was through a dual-pass 420-to 480- and 570-to 620-nm Chroma filter, while 561-nm was collected through the dual-pass filter or the specific RED 596-to 696-nm Chroma filter, depending on the experiment. The 640-nm excitation wavelength was collected by 505 to 550-nm and 650 to 850-nm dual pass Chroma filter, while 488 nm was collected through the dual-pass filter or the specific FITC 490 to 525-nm Chroma filter, again depending on the experiment and degree of crosstalk among fluorophores in the sample.

### Data Analysis

All experiments conducted with confocal and TIRF assemblies produced nd2 files that were opened and analyzed through ImageJ implementation, Fiji^62^.

### Image segmentation to obtain organelle-specific masks

Nd2 files were imported into FIJI and analyzed by custom macros^63^. ROIs were drawn with the Freehand selection tool around the footprint of whole cells. Organelle-specific masks were generated by filtering images of the marker-containing channel with a Gaussian blur filter set as multiples of the point spread function (PSF) of the marker’s fluorophore. The filtered images were used to generate wavelets by subtracting each channel’s image by the next smallest filter. The wavelet standard deviation (SD) was then calculated and multiplied by 0.5 to set a hard threshold. The resulting wavelet products were calculated by multiplying the wavelets together, before converting to a binary Mask^64^.

### Single molecule analysis

Single-molecule imaging was performed using TIRF microscopy, which allowed measurement of the acutely expressed biosensors and endogenous NeonGreen tagged particles at the cell surface within a defined 100μm^2^ region of interest. Fiji plugin ThunderSTORM^65^ was used to count single molecules. For single particle colocalization estimations, the molecules are localized with approximately 20nm precision, and organelle markers segmented as described in the prior section. Particle density was then calculated by counting all particles in the 100μm^2^ region of interest, or just those localizing within the segmented organelle masks. Both densities were converted to particles per 100μm^2^. To calculate the percentage of particles at a specific organelle marker, the particle count co-localized with the segmented image was divided by the total particle count across the whole region of interest.

### z-stack Imaging

To image the z-stacks of cells, the Nikon A1 Piezo Z Drive was employed to rapidly acquire time-lapse images of the entire volume of live-cells. The stack imaging scanned from bottom to top of the cell within the 15μm stack with 0.175 μm steps (approximately Nyquist sampling). An average z projection between steps 40-50 were used to both analyze and present the data as this centered on the middle of the cell where the plasma membrane and endosomes could be best visualized. The biosensor localization to the specific structures was represented by ratio measurement Organelle/Cell.

### PM intensity changes by TIRFM

To measure PM fluorescence intensity, ROIs were drawn with the Freehand selection tool around a minimum intensity projection of the cell footprint. The minimum projection accounts for cell movement during timelapse imaging, however cells with excessive movement were precluded from analysis. The images were imported into FIJI and analyzed with custom-written macros that measured the fluorescence intensity within the set ROI at each frame in the post-stimulation time lapse (F_t_), normalized to the baseline pre-stimulation frames (F_pre_).

### Fluorescence Intensity

To acquire confocal images of weakly expressed, endogenously tagged mNeonGreen protein, we acquired images from both green (500-550 nm) and orange/red (570-620 nm) channels using 488 nm excitation. Cellular autofluorescence present in both 488 and 561 channel was first removed by subtracting the red from the green channel. For confirmation of enzyme-specific mNeonGreen integration, the cell lines were genotyped by comparison of cellular fluorescence intensity measured in the unedited-parental HEK293A cell line to the edited lines treated with non-targeting control and specific siRNA (Figure 6A-B). ROIs were drawn with the Freehand selection tool around whole cell footprints in the DIC channel. The fluorescence intensity within the ROIs were then measured in the corrected 488-projection by FIJI and could be compared between control and siRNA treated cells.

### Key resources table

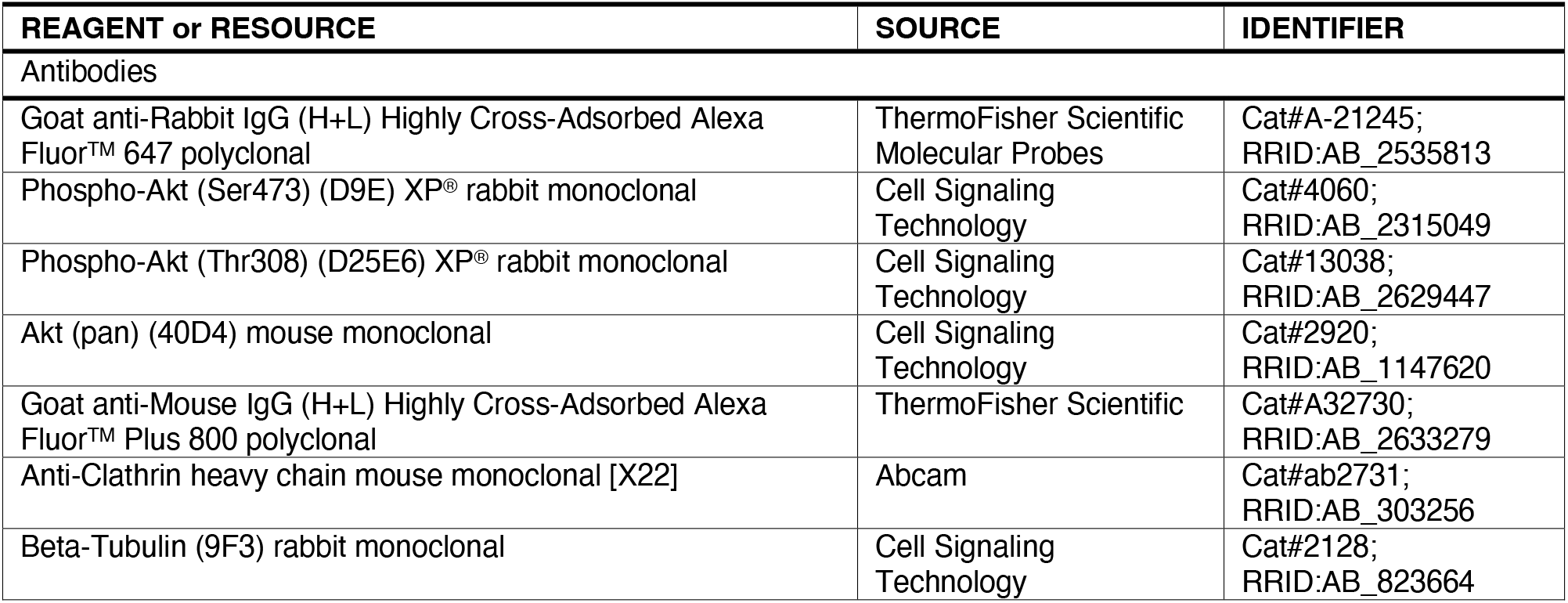
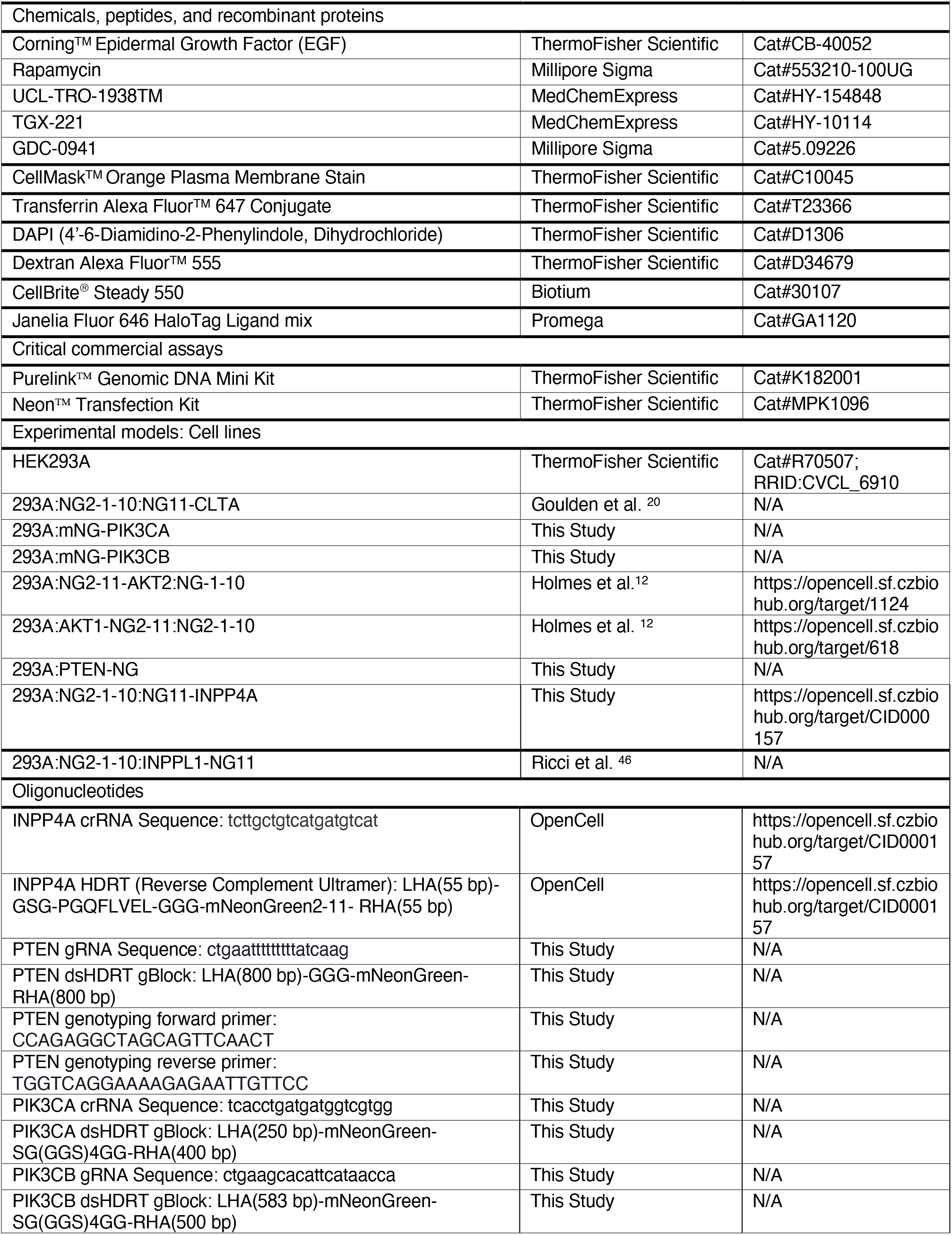
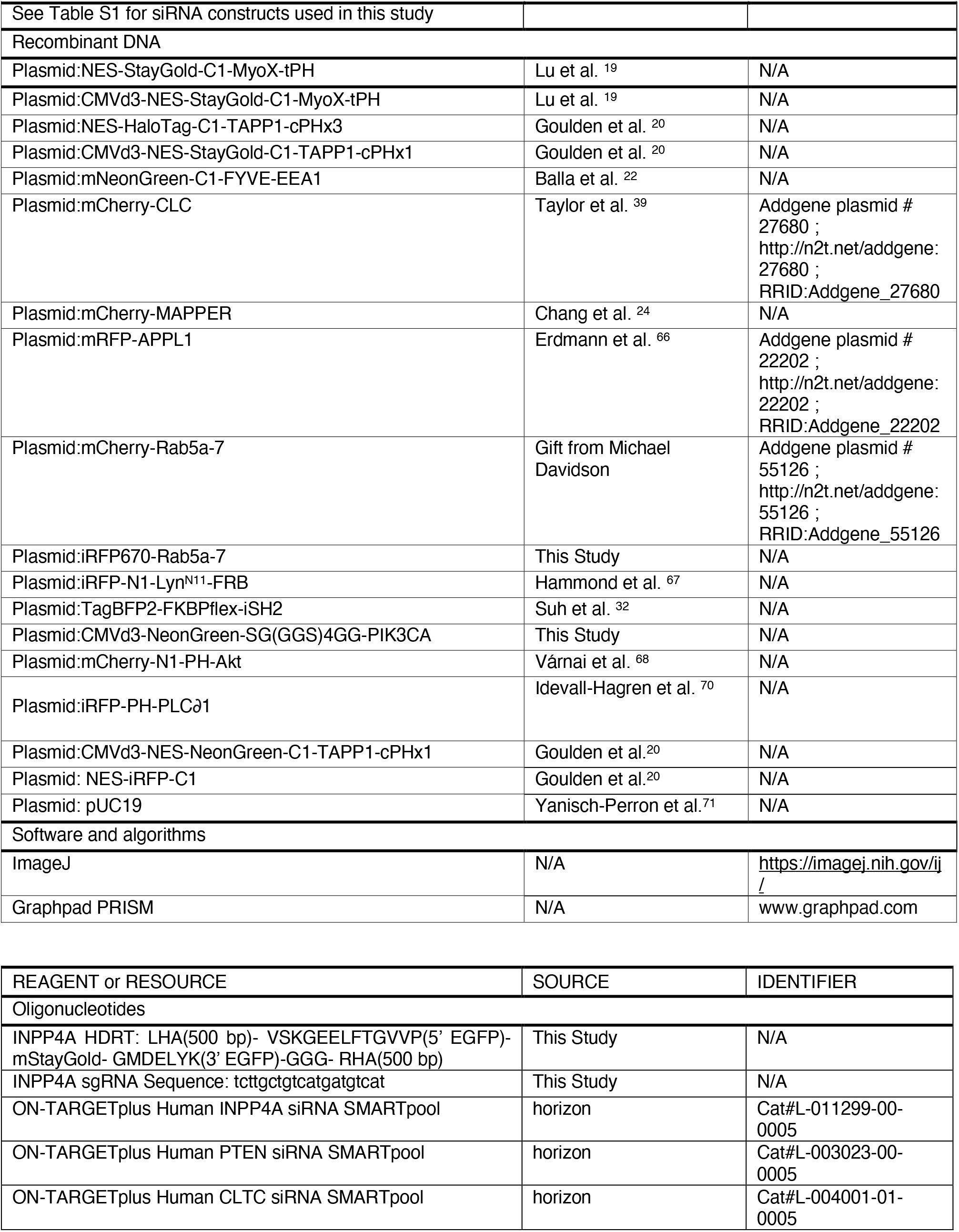

## Author Contributions

Morgan M. C. Ricci: Investigation; Visualization; Writing - Original Draft; Writing - Review & Editing.

Hemani Patel, Maria Montoya, Kanishka Jayasuriya, Andrew Rectenwald and Magdelene A. K. Motter: Investigation.

Gerald R.V. Hammond: Conceptualization; Data analysis; Writing - Original Draft; Writing - Review & Editing.

---

**FIGURE S1:**
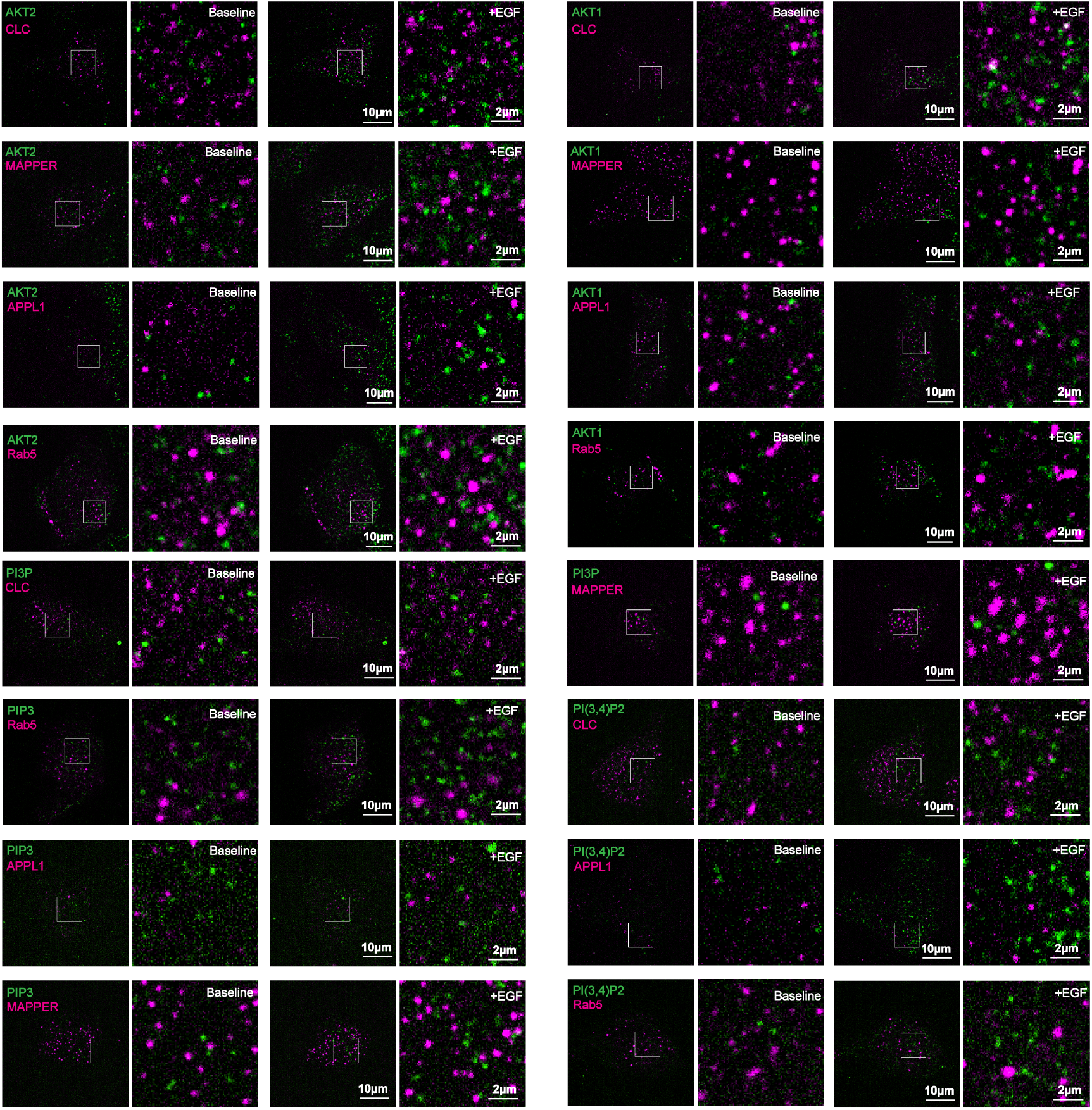
Representative images of endogenous NG2-tagged AKT1 or AKT2 or the indicated lipid biosensors, together with organelle markers pre- and post-EGF administration. For each image pair, the left image shows then entire cell footprint, whereas the right image shows the inset.

**FIGURE S2:**
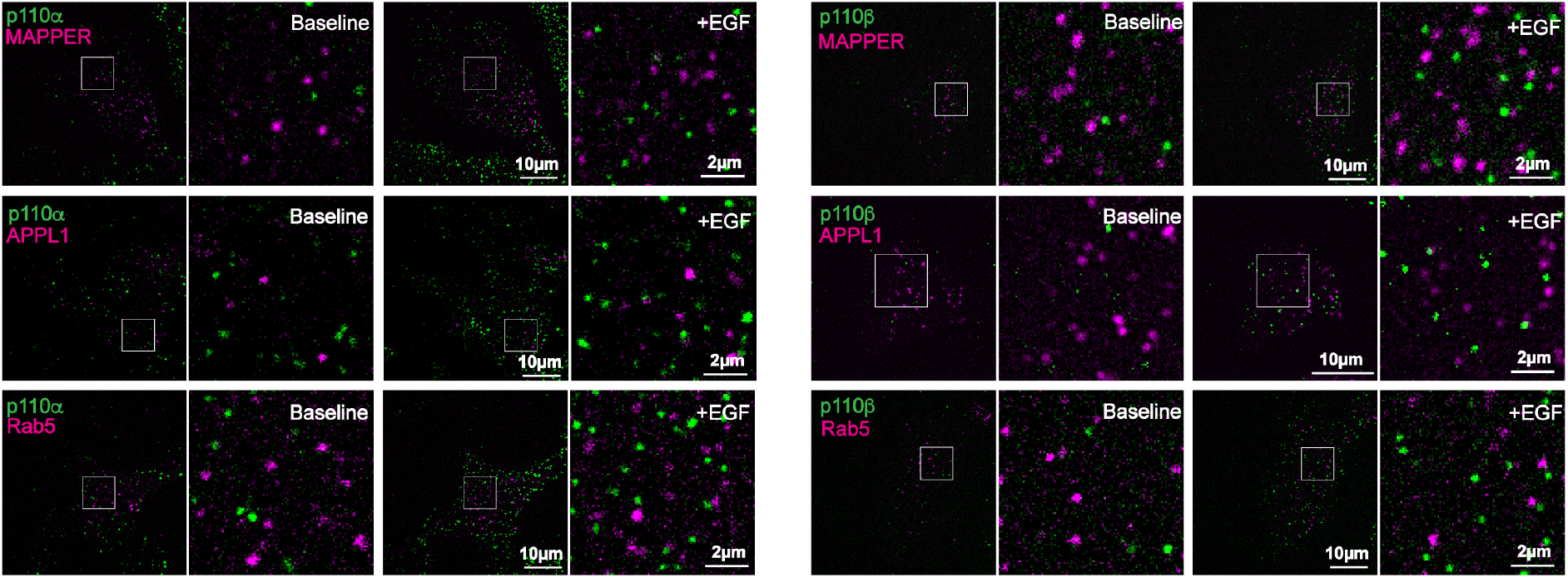
Representative images of endogenous mNeonGreen-tagged p110α or p110β, together with organelle markers pre- and post-EGF administration. For each image pair, the left image shows then entire cell footprint, whereas the right image shows the inset.

**FIGURE S3:**
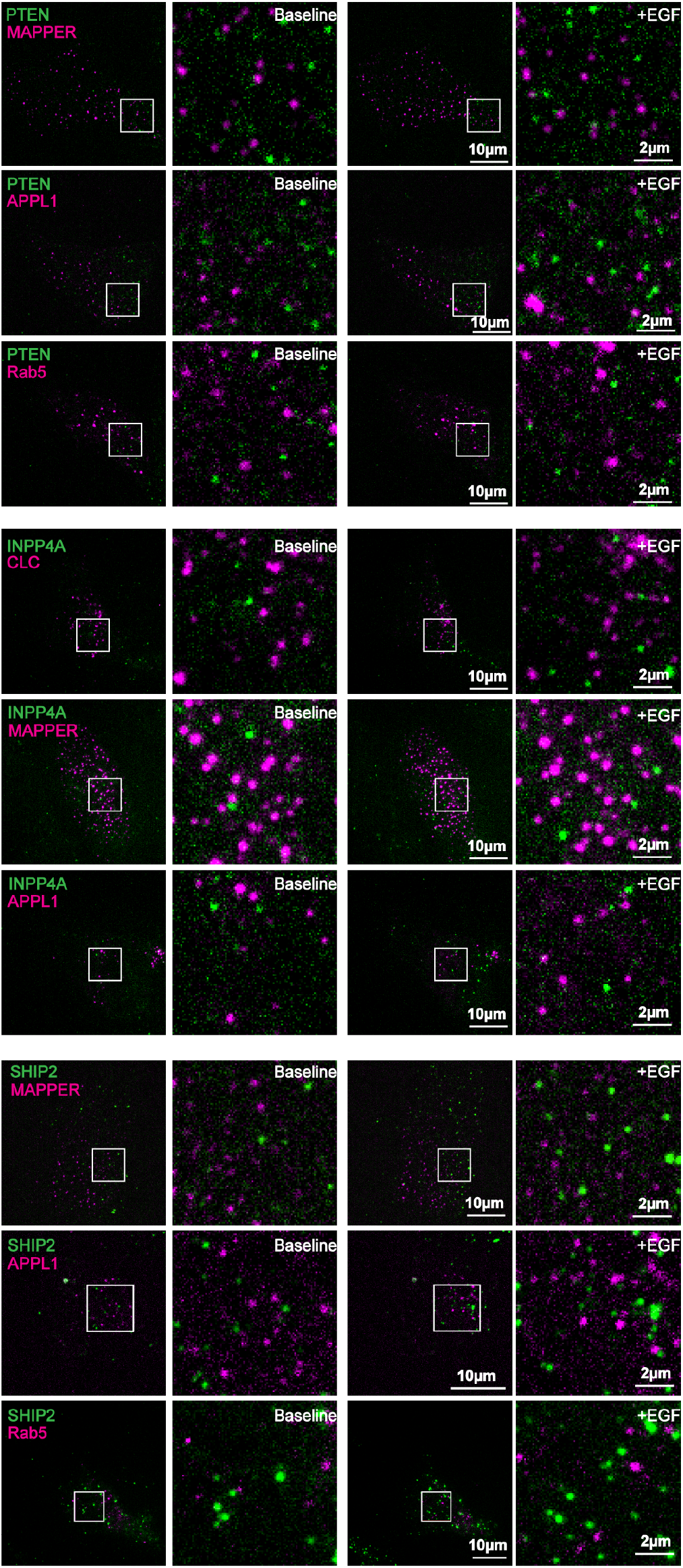
Representative images of endogenous tagged PTEN, INPP4A or SHIP2, together with organelle markers pre- and post-EGF administration. For each image pair, the left image shows then entire cell footprint, whereas the right image shows the inset.

